# GR-SAFS: A Graph-Regularized Stacking Framework with Adaptive Feature Selection for High-Dimensional Prognostic Biomarker Discovery

**DOI:** 10.64898/2026.06.23.733986

**Authors:** Jin-Zhao He, Jing Guan

**Author notes:** Corresponding author: Jing Guan.

## Abstract

Identifying prognostic biomarkers from high-dimensional transcriptomic data poses a triple challenge: achieving sparsity, preserving biological network topology, and integrating complementary nonlinear signals. Existing methods typically ignore network structure, miss nonlinear interactions, or lack a principled mechanism to fuse heterogeneous model outputs. We introduce GR-SAFS (Graph-Regularized Stacking with Adaptive Feature Selection), a framework with three modules: a Graph-Lasso engine embedding gene co-expression network Laplacian priors, run in parallel with a Random Forest engine; an empirical cumulative distribution function (eCDF) alignment layer that places sparse and dense importances on a common percentile scale; and a diversity-penalized quadratic programming router whose strict convexity yields a unique global optimum. On the TCGA-LUAD cohort, GR-SAFS identifies a 20-gene signature with a training concordance index of 0.700. Across two independent cross-platform microarray cohorts, GR-SAFS is the only method whose frozen signature retains statistically significant risk stratification in every cohort, where stronger-C-index baselines lose significance on at least one external cohort. Functional enrichment anchors the signature to a coherent Wnt/*β*-catenin axis. An open-source implementation is released for full reproducibility.

## I. Introduction

**L**UNG adenocarcinoma (LUAD) is the most common histological subtype of lung cancer, accounting for 40–50% of all cases. Despite advances in targeted therapy, substantial interpatient variability in five-year survival persists even within the identical TNM stage [1]. The discovery of molecular prognostic biomarkers based on gene expression profiling has consequently emerged as a critical research direction in precision oncology [2].

High-throughput sequencing has rendered whole-transcriptome profiling routine, yet it introduces the classical high-dimensional small-sample (*p* ≫*n*) challenge: typical transcriptomic data contain approximately 10^4^ genes while clinical cohorts provide only 10^2^–10^3^ samples, with the truly prognosis-related genes constituting a minimal fraction of the candidates. Existing high-dimensional feature selection methods encounter a triple contradiction. Linear *ℓ*_1_-regularized methods such as Lasso [3] and Elastic Net [4] offer theoretical guarantees but ignore biological associations among genes. This limitation results in module fragmentation in the presence of co-linear gene clusters [4], which subsequently leads to poor reproducibility and the loss of biological interpretability.

Nonparametric machine learning methods such as Random Forest [5] and Random Survival Forest [6] implicitly capture nonlinear interactions but produce dense, non-sparse importance scores, and their tree-based split gains are incommensurable with linear regression coefficients, thereby creating a metric gap that precludes direct fusion. Recent reviews have systematically evaluated deep learning approaches for survival analysis [7], highlighting both the expressiveness of these approaches and the persistent challenge of overfitting in high-dimensional regimes. Recent multiply robust estimation methods for partially linear models [8] achieve robustness across data conditions but do not exploit biological network topology, limiting their applicability to transcriptomic-scale prognostic modeling.

Network-constrained regularization methods, including Network-Constrained Lasso [9] and dwgLASSO [10] incorporate biological topology priors but typically employ single linear engines, which fail to capture higher-order nonlinear interactions. Beyond purely linear engines, graph-spectral priors have also been combined with nonparametric models, for example Gaussian processes with spectral graph features for graph-level classification [11]; such combinations, however, have not been developed for high-dimensional prognostic feature selection.

To unify these three requirements, this study proposes GR-SAFS. The primary contributions are as follows:

1. For LUAD prognostic biomarker discovery, we couple gene co-expression network Laplacian priors with a nonparametric ensemble engine within a single selection pipeline, so that topology-aware linear main effects and nonlinear interactions are captured together rather than by separate models.
2. We introduce an empirical-CDF alignment layer that places the sparse Graph-Lasso coefficients and the dense ensemble importances on a common percentile scale, removing the metric gap that prevents their direct fusion.
3. We formulate adaptive weight allocation as a diversity-penalized quadratic program whose strict convexity yields a unique global optimum (Proposition 1), and we show empirically that the diversity penalty curbs cross-platform overfitting.
4. Empirical results on TCGA-LUAD and two independent cross-platform validation cohorts show the cross-platform generalization robustness of GR-SAFS; the resulting 20-gene signature is biologically coherent, with all top-10 candidates supported by independent LUAD literature and a coordinated Wnt/*β*-catenin axis recovered through multi-ontology enrichment. A reproducible open-source implementation is released (Section V).

## II. Materials and Methods

### A. Data and Preprocessing

This study utilized three LUAD cohorts. The training cohort was TCGA-LUAD (504 patients with valid survival data) using RNA-seq profiles from UCSC Xena [12]. The two external validation cohorts were GSE31210 (Japan National Cancer Center, *n* = 226, stage I–II) [13] and GSE50081 (Canadian multi-centers, *n* = 181) [14], both based on Affymetrix HG-U133 Plus 2.0 microarrays. Clinical survival data were obtained from TCGA-CDR [15].

The preprocessing steps included: (1) mapping Ensembl IDs to HGNC Gene Symbols via MyGene.info; (2) removing duplicates while retaining the highest-mean transcript; (3) applying a log_2_(FPKM + 1) transformation; and (4) retaining the top 10,000 most-variable genes. Among the top 20 candidate genes of GR-SAFS, 19 genes were successfully matched in each validation cohort. To prevent inflated generalization estimates, a strict frozen signature paradigm was adopted: the 20-gene list, the effect directions, and the fusion weights determined on TCGA-LUAD were held constant on the validation sets.

### B. Dual-Core Feature Scanning

As illustrated in Fig. 1, GR-SAFS deploys two heterogeneous engines in parallel on the full *p*-dimensional feature space. **Knowledge-driven Graph-Lasso engine**. Given the standardized feature matrix **G** ∈ℝ^*n×p*^ and the phenotype **y** ∈ℝ^*n*^, a sparse adjacency matrix **A** is first constructed from the Pearson correlation matrix **R** via soft thresholding (threshold *τ* = 0.3):

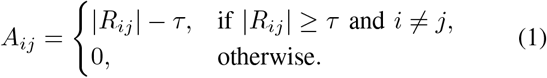

**Fig. 1:**
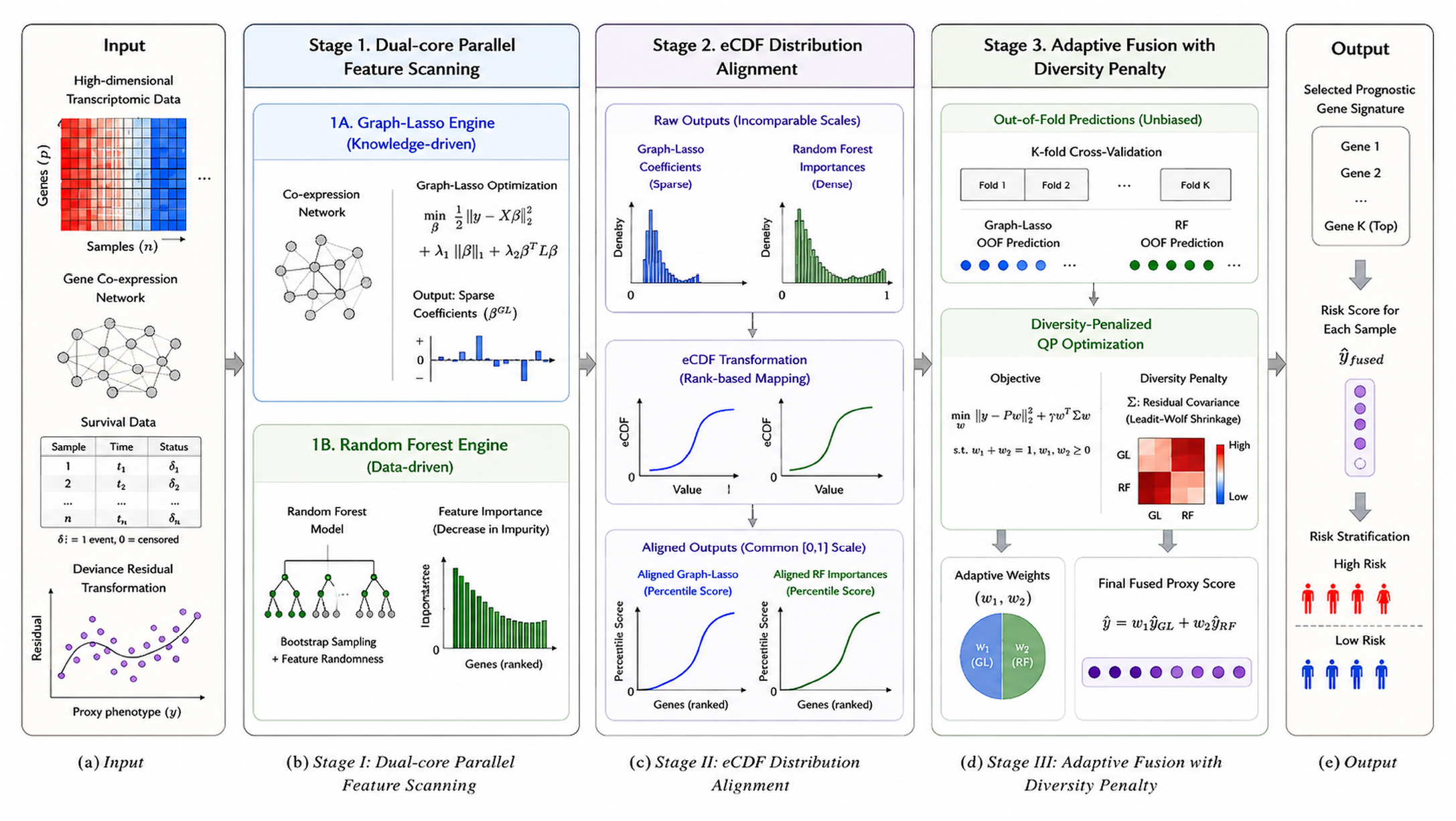
The GR-SAFS framework. (a) Stage 1: Dual-core parallel scanning deploys a knowledge-driven Graph-Lasso engine (embedding the gene co-expression network Laplacian **L**) and a data-driven Random Forest engine on the full *p*-dimensional feature space. (b) Stage 2: eCDF distribution alignment maps heterogeneous outputs—sparse Graph-Lasso coefficients and dense RF importance scores—to a unified percentile space [0, 1]. (c) Stage 3: Adaptive fusion via diversity-penalized quadratic programming optimizes weights through out-of-fold cross-predictions, with the Ledoit–Wolf shrinkage of the residual covariance matrix forcing error decorrelation.

The graph LapI:lacian **L** = **D** − **A** is subsequently derived with *D*_*ii*_ = *Σ*_*j*_ *A*_*ij*_. The threshold *τ* = 0.3 is not a sensitive hyperparameter: across *τ* ∈ [0.20, 0.40] the training C-index varies by less than 0.01 and at least 17 of the 20 signature genes are retained, with risk stratification remaining significant throughout (Supplementary Material, Section S6.2). The generalized graph-regularized objective is formulated as:

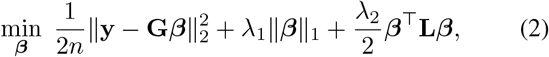

where 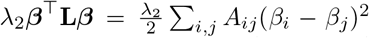 enforces smooth coefficients on topologically adjacent nodes. In a Bayesian interpretation, this formulation corresponds to a multivariate Laplacian prior on ***β*** that depends on the graph structure. This non-smooth problem is solved via proximal gradient descent (PGD) with a fixed step size equal to the inverse Lipschitz constant of the smooth gradient (computed once as the largest Hessian eigenvalue by a single Lanczos iteration); the solver has an *O*(1*/k*) convergence rate and converges on the full *p* = 10,000 feature space in under 40 seconds on a single CPU core (Supplementary Material, Section S2).

### Unified survival-to-phenotype transformation

To process right-censored survival data within the MSE-based Graph-Lasso optimization pipeline, we apply a deviance-residual transformation under the null Cox model (i.e., no covariates), following Therneau et al. [16]. Given observation time *t*_*i*_ and event indicator *δ*_*i*_ ∈ {0, 1}, the Nelson–Aalen estimator of the cumulative baseline hazard is 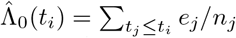, where *e*_*j*_ denotes the number of events at distinct failure time *t*_*j*_ and *n*_*j*_ the number of subjects at risk just prior to *t*_*j*_. The martingale residual under the null model is 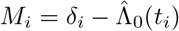, and the deviance residual is obtained by the variance-stabilizing transformation

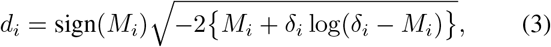

with the convention *δ*_*i*_ log(*δ*_*i*_ − *M*_*i*_) := 0 when *δ*_*i*_ = 0. Because the Graph-Lasso engine minimizes a least-squares objective, the deviance residual only needs to be a well-behaved, finite-variance continuous surrogate rather than a strictly normal variable [16]; we use it as **y** in (2). A detailed derivation and an empirical analysis of its distribution on TCGA-LUAD (variance-stabilized but moderately right-skewed) are provided in the Supplementary Material, Section S1.3.

### Data-driven Random Forest engine

In parallel, a regression Random Forest (300 trees, maximum depth 5, minimum leaf size 3) is trained on the continuous deviance-residual surrogate **y** and scans the full feature space via recursive tree splits. Because the target is continuous, node impurity is measured by the within-node variance (mean squared error) of **y**, not by a classification criterion. For a node *v* that is split on feature *x*_*j*_, let ΔVar(*v, x*_*j*_) denote the resulting weighted decrease in target variance. The nonlinear importance score is the total variance reduction attributable to feature *j*, averaged over the forest:

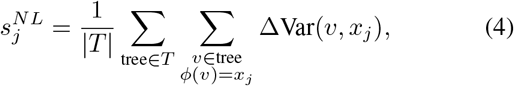

which outputs a dense vector 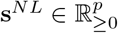. Training a regression forest on the deviance residual, rather than a classifier on event status, ensures that the importance reflects the full survival information encoded in **y** (events and censoring, through the martingale transformation) rather than a dichotomized outcome.

### C. eCDF Distribution Alignment

To eliminate the metric gap between sparse Graph-Lasso coefficients (extreme values at few genes, and zeros elsewhere) and dense Random Forest scores, a percentile mapping based on ascending ranks is introduced. For any importance vector 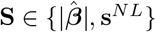, let rank_min_(*S*_*i*_) be the ascending rank of *S*_*i*_, with tied values assigned their smallest rank. The mapped percentile scores are

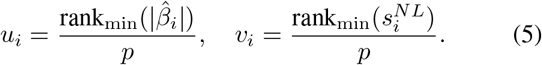

This converts absolute magnitudes into relative ranks, representing the proportion of background features that a given gene surpasses in the respective engine. The min-rank convention is essential for the sparse Graph-Lasso output: the zero coefficients form the smallest and most numerous tied block, so they all receive the smallest rank and are mapped to the percentile floor (*u*_*i*_ = 1*/p* when 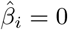), which truncates them as uninformative. By contrast, the ordinary empirical CDF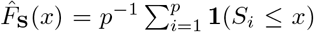, which counts all ties, would map every zero to ≈ *p*_0_*/p* (where *p*_0_ is the number of zeros), placing them near the top; (5) avoids this and is the mapping used throughout. These mappings inherit the standard properties of the empirical CDF (distribution invariance, rank preservation, and monotonic invariance), stated in the Supplementary Material (Propositions S1–S3) for completeness; they are elementary properties that hold for any monotone transformation and are not claimed as a theoretical contribution.

### D. Diversity-Penalized QP Fusion

The stacking ensemble paradigm [17], originally proposed by Wolpert, has been widely adopted in modern ensemble learning [18]. The proposed QP routing mechanism extends this paradigm by introducing diversity-aware regularization. Through *K* = 5 out-of-fold (OOF) cross-predictions, the metaprediction matrix 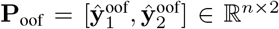 is constructed,and the weight allocation is formulated as:

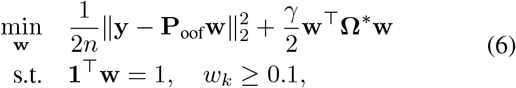

where **Ω**^*\**^ represents the Ledoit–Wolf [19] linearly shrunk residual covariance matrix. The diversity penalty **w**^*⊤*^**Ω**^*\**^**w** is equivalent to the joint prediction variance of the ensemble, forcing error-decorrelated model selection. The lower bound *w*_*k*_ ≥ 0.1 ensures a substantive contribution from both signal sources.

#### Proposition 1

(Existence and Uniqueness of the QP Solution). *Consider the weight-allocation problem in* (6) *over m base engines, with weight vector* **w** ∈ℝ^*m*^ *and feasible set* = **w** ∈ℝ^*m*^ : **1**^*⊤*^**w** = 1, *w*_*k*_ ≥ *ℓ for k* = 1, …, *m, where ℓ* ∈ [0, 1*/m*] *is the per-engine lower bound (m* = 2 *and ℓ* = 0.1 *in GR-SAFS). Let* 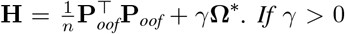 *and* **Ω**^*\**^ *is positive definite after the Ledoit–Wolf shrinkage, then* C *is non-empty, compact, and convex, the objective is strictly convex, and* (6) *admits a unique global minimizer* **w**^*\**^ ∈ *C*.

*Proof. Feasibility and compactness*. The uniform vector 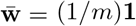 satisfies 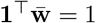 and ≥ *ℓ* since *ℓ* ≤ 1*/m*; hence 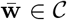 and *C* ≠ ∅. The set C is the intersection of the hyperplane {**1**^*⊤*^**w** = 1} and the box {*w*_*k*_ ≥ *ℓ*, ∀*k*}, both closed and convex, so C is closed and convex. It is bounded, because for every 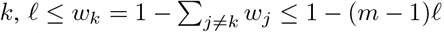 By the Heine–Borel theorem, C is therefore compact.

*Strict convexity*. With 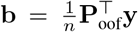, the objective of (6) equals 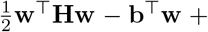 const, whose Hessian is **H**. For any **v**≠ **0** in 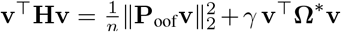. The first term is non-negative; the second is strictly positive because **Ω**^*\**^ ≻ 0 after the Ledoit–Wolf shrinkage and *γ >* 0. Hence **H** ≻ 0, and the objective is strictly convex on ℝ^*m*^, and a fortiori on the convex set *C*.

*Existence and uniqueness*. The objective is continuous and is non-empty and compact, so by the Weierstrass extreme value theorem a global minimizer exists on. A strictly convex function admits at most one minimizer over a convex set [20]; therefore the global minimizer **w**^*\**^ ∈ *C* is unique.

#### Proposition 2

(Stability of the Fusion Weights). *Under the assumptions of Proposition 1, write the data-coupling vector as* 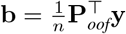, *so that* **w**^*⋆*^(**b**) = arg min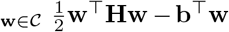.*Let µ* = *λ*_min_(**H**) ≥ *γ λ*_min_(**Ω**^*\**^) *>* 0. *Then for any two coupling vectors* **b**_1_, **b**_2_ *with optimal weights*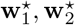,

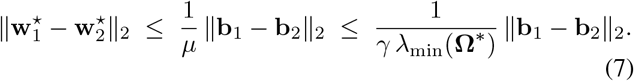

*That is, the fusion weights depend Lipschitz-continuously on the data, with a modulus controlled by the diversity strength γ and the smallest shrunk-covariance eigenvalue λ*_min_(**Ω**^*\**^).

*Proof*. By Proposition 1, **H** ≻ 0, so the objective is *µ*-strongly convex with 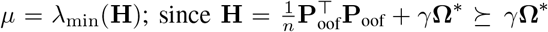, we have *µ* ≥ *γ λ*_min_(**Ω**^*\**^). Each minimizer over the closed convex set C satisfies the first-order variational inequality 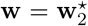 for all **w** ∈ C. Taking 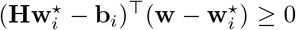 in the inequality for *i* = 1 and 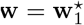 for *i* = 2 and adding,

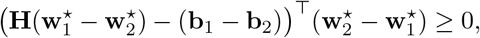

which rearranges to 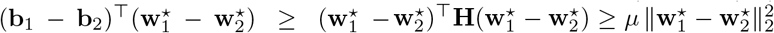. Applying the Cauchy– Schwarz inequality to the left-hand side and dividing by 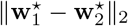 (the bound is trivial when the weights coincide) yields (7).

Proposition 2 explains, at the level of the fusion step, why the Ledoit–Wolf shrinkage is not merely a numerical safeguard: by lifting *λ*_min_(**Ω**^*\**^) away from zero it bounds the sensitivity of the routing weights to data perturbations, which in turn underpins the high selection reproducibility reported in Section III. A non-shrunk, possibly singular residual covariance would leave this modulus unbounded.

The feature-level fusion score is calculated as:

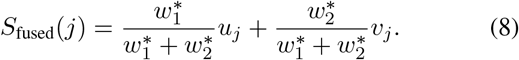

The top-*K* candidates are selected in descending order of *S*_fused_. The QP is a two-variable convex program solved in milliseconds; variance-reduction analysis and a permutation-importance validation of the weight transfer are provided in the Supplementary Material, Section S5.

### E. Experimental Setup

Recent benchmark studies regarding feature selection for survival data [21] have highlighted that no single method demonstrates dominance across all scenarios, which motivates the heterogeneous fusion approach employed in this study.

### Rationale for the simulation design

We evaluate feature-selection accuracy on simulated outcomes because clinical cohorts provide no ground-truth set of causal genes. Recovery metrics such as TPR, FPR, and AUPRC require a known generative model, which real transcriptomes cannot supply; the real cohorts therefore serve to test prognostic generalization through the C-index, the log-rank test, and frozen-signature external validation, rather than to measure selection accuracy. Crucially, the simulation does not impose a synthetic block-diagonal correlation structure. The feature matrix consists of real human genotypes from the 1000 Genomes Project, so the inter-feature correlation is the empirical linkage-disequilibrium structure of thousands of real genomes; only the phenotype is simulated, by injecting known causal variants, pleiotropy, and epistasis (Supplementary Material, Section S7). This yields a realistic and intricate correlation pattern with a known causal ground truth, which a block-diagonal toy model could not reproduce.

### Hyperparameter optimization

A nested cross-validation (outer 3-fold× inner 3-fold) with a two-stage decoupled search was adopted: stage 1 optimizes (*λ*_1_, *λ*_2_) by minimizing the inner MSE; stage 2 optimizes *γ* by minimizing the OOF residuals. On TCGA-LUAD, the optimal configuration was 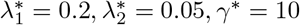.

### Baseline methods

For the simulation benchmark, GR-SAFS was compared against nine representative high-dimensional feature selection methods spanning linear shrinkage (Univariate, Lasso [3], Elastic Net [4]), structured sparsity (Group Lasso [22], Sparse Group Lasso [23], Graph Lasso [9], Precision Lasso [24]), controlled-error inference (Knockoffs [25]), representation learning (VAE [26]), and nonparametric ensembles (Random Forest [5]). For the real-data prognostic comparison, seven survival-specific baselines were employed, covering linear regularized models (Cox-Lasso [27], Cox-Elastic Net [4], Univariate Cox, NC-Lasso [9]) and modern nonlinear/deep survival models (CoxBoost [28], RSF [6], DeepSurv [29]). Each method performed its own end-to-end feature selection down to an identical 20-feature budget for a fair comparison; complete hyperparameters are listed in the Supplementary Material, Section S9. Filter-based selectors such as mRMR, widely used in moderate-dimensional clinical prediction [30], were not included as they target mutual-information ranking rather than *p* ≫ *n* sparse regression.

### Evaluation metrics

The evaluation used the C-index for ranking ability, the Kaplan–Meier log-rank test for stratification significance, the 1-year AUC for short-term discrimination, F1/Precision/Recall for classification under the median-risk threshold, and the AUPRC with the median rank for simulation accuracy and stability.

## III. Results

### A. Simulation Validates Cross-Architecture Robustness

Three progressively challenging scenarios were constructed: S1 (additive: 10 causal variants, *h*^2^ = 0.7), S2 (network pleiotropy: plus 20 weak pleiotropic nodes, *h*^2^ = 0.5, epistasis 5.0), and S3 (extreme epistasis: *h*^2^ = 0.4, epistasis 10.0). Under the condition of *p* = 10,000, GR-SAFS exhibited robust cross-scenario balance (Table I, Fig. 2). In the most challenging S3 scenario, GR-SAFS matched the AUPRC of the Random Forest model (0.337 vs. 0.337) while achieving a substantially superior median rank (1027 vs. 2029), which demonstrates that graph-regularized priors effectively constrain the search space. In the additive S1 scenario, where no co-expression structure links the causal variants, GR-SAFS attained a lower AUPRC (0.784) than the pure sparse regressors Lasso (0.904), Precision Lasso (0.906), and Knockoffs (0.903); its advantage over these methods emerged only as nonlinear interaction and pleiotropy increased. Across the three scenarios, the relative AUPRC of GR-SAFS rose monotonically from 86.6% of the per-scenario maximum in S1 to 100% in S3 (Fig. 2c), the only method to show this pattern.

**TABLE I:** Comprehensive simulation benchmark at *p* = 10,000 across three genetic architectures. All values are averaged over 20 independent simulation runs. The optimal value in each column is in bold. S1: additive baseline, 10 causal variants, *h*^2^ = 0.7, no nonlinear interactions. S2: network pleiotropy, 20 weak pleiotropic nodes, *h*^2^ = 0.5, epistasis 5.0. S3: extreme epistasis, *h*^2^ = 0.4, epistasis 10.0.

| Scenario | Model | TPR | AUPRC | FPR | Rank | Precision |
| --- | --- | --- | --- | --- | --- | --- |
| S1 | GR-SAFS | 0.760 | 0.784 | 2.40E-04 | 93.7 | 0.760 |
|  | Univariate | 0.500 | 0.511 | 5.01E-04 | 525.9 | 0.500 |
|  | Lasso | **0.880** | 0.904 | **1.20E-04** | **5.6** | **0.880** |
|  | ElasticNet | 0.855 | 0.882 | 1.45E-04 | 5.6 | 0.855 |
|  | GroupLasso | 0.640 | 0.664 | 3.60E-04 | 12.8 | 0.640 |
|  | GraphLasso | 0.780 | 0.807 | 2.20E-04 | 5.9 | 0.780 |
|  | PrecisionLasso | 0.880 | **0.906** | 1.20E-04 | 5.6 | 0.880 |
|  | Knockoffs | 0.880 | 0.903 | 1.20E-04 | 5.6 | 0.880 |
|  | VAE | 0.505 | 0.525 | 4.95E-04 | 485.3 | 0.505 |
|  | RF | 0.725 | 0.749 | 2.75E-04 | 35.6 | 0.725 |
| S2 | GR-SAFS | **0.435** | 0.376 | **5.66E-04** | 555.3 | **0.435** |
|  | Univariate | 0.330 | 0.282 | 6.71E-04 | 1114.3 | 0.330 |
|  | Lasso | 0.425 | 0.371 | 5.76E-04 | **186.6** | 0.425 |
|  | ElasticNet | 0.425 | 0.356 | 5.76E-04 | 245.4 | 0.425 |
|  | GroupLasso | 0.365 | 0.318 | 6.36E-04 | 325.1 | 0.365 |
|  | GraphLasso | 0.410 | 0.367 | 5.91E-04 | 974.4 | 0.410 |
|  | PrecisionLasso | 0.425 | 0.371 | 5.76E-04 | 195.9 | 0.425 |
|  | Knockoffs | 0.430 | **0.382** | 5.71E-04 | 327.9 | 0.430 |
|  | VAE | 0.285 | 0.226 | 7.16E-04 | 1304.6 | 0.285 |
|  | RF | 0.395 | 0.344 | 6.06E-04 | 1027.5 | 0.395 |
| S3 | GR-SAFS | **0.375** | **0.337** | **6.26E-04** | **1026.8** | **0.375** |
|  | Univariate | 0.290 | 0.254 | 7.11E-04 | 1767.1 | 0.290 |
|  | Lasso | 0.355 | 0.315 | 6.46E-04 | 1318.1 | 0.355 |
|  | ElasticNet | 0.355 | 0.304 | 6.46E-04 | 1250.2 | 0.355 |
|  | GroupLasso | 0.320 | 0.277 | 6.81E-04 | 1168.8 | 0.320 |
|  | GraphLasso | 0.355 | 0.316 | 6.46E-04 | 1846.7 | 0.355 |
|  | PrecisionLasso | 0.370 | 0.317 | 6.31E-04 | 1312.7 | 0.370 |
|  | Knockoffs | 0.365 | 0.315 | 6.36E-04 | 1055.1 | 0.365 |
|  | VAE | 0.245 | 0.215 | 7.56E-04 | 1960.6 | 0.245 |
|  | RF | 0.365 | 0.337 | 6.36E-04 | 2029.3 | 0.365 |

**Fig. 2:**
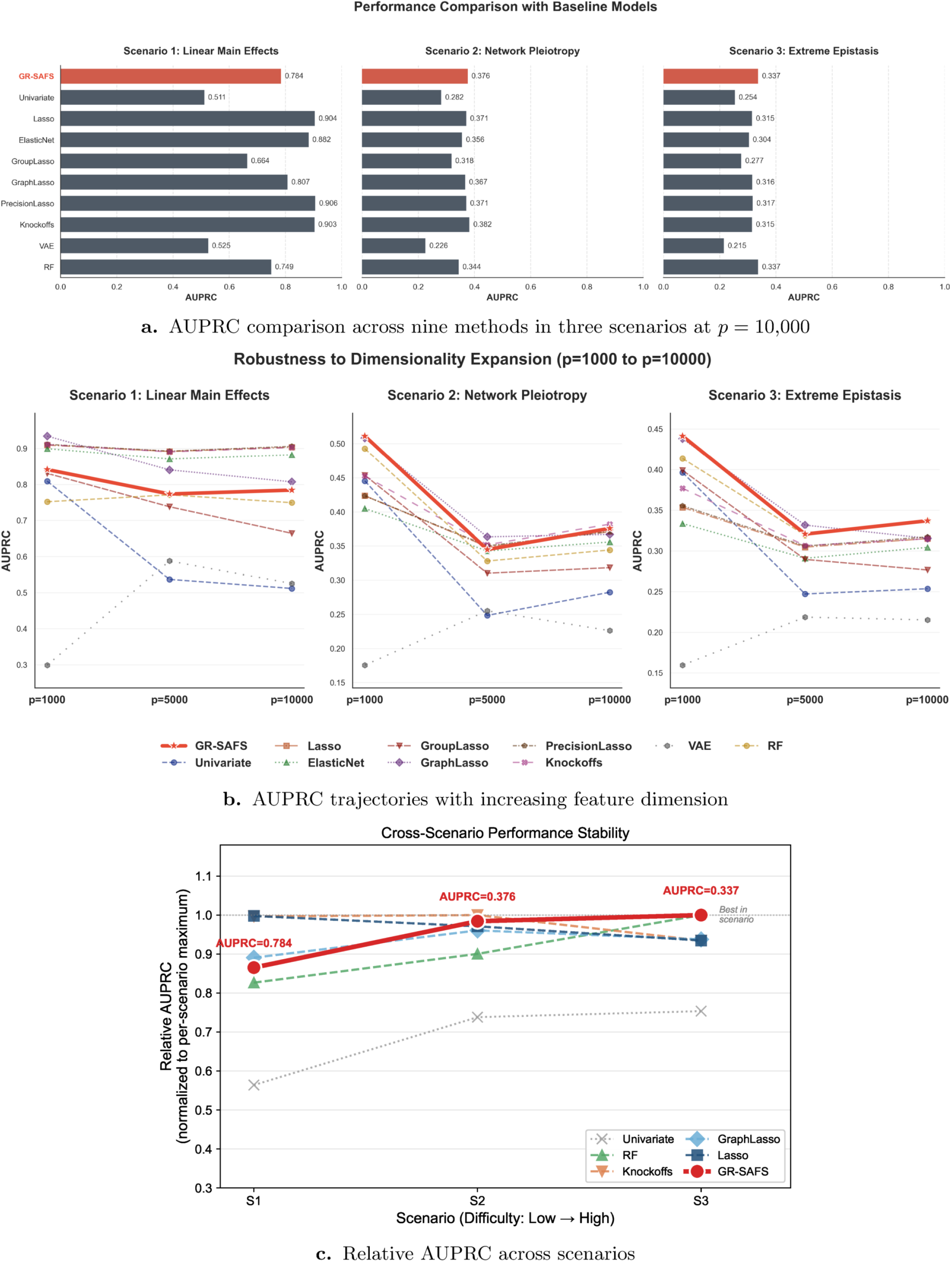
Simulation benchmark. (a) AUPRC comparison across nine methods in three scenarios (S1 additive, S2 network pleiotropy, S3 extreme epistasis) at *p* = 10,000. (b) AUPRC trajectories with increasing feature dimension (*p* = 1,000, 5,000, 10,000). GR-SAFS stabilizes beyond *p* = 5,000 while traditional methods decline monotonically. (c) Relative AUPRC normalized to the per-scenario maximum. GR-SAFS demonstrates the only monotonically increasing trajectory, rising from 86.6% in S1 to 100% in S3.

Dimension-scaling experiments (*p* ranging from 1,000 to 10,000) further highlighted the robustness of the framework (Fig. 2b). Traditional methods (Univariate, Lasso) exhibited a monotonic decline in AUPRC as the dimension increased. Conversely, GR-SAFS stabilized beyond *p* = 5,000 and demonstrated a slight upward trend in S3. We report this stabilization as an empirical observation rather than a formal effect. It is consistent with the diversity penalty promoting error cancellation as the number of noise features grows, although we do not claim a theoretical guarantee for it.

### B. Ablation Study of GR-SAFS Components

To dissect the contribution of each module, four variants were constructed: GL only (knowledge-driven solely), Uniform 1/2 (equal-weight fusion), MSE Stack (QP with *γ* = 0, strictly accuracy-driven), and the complete GR-SAFS framework (Fig. 3). By leveraging strong linear priors, GL only achieved the highest AUPRC in S1 (0.823), yet collapsed to 0.301 in S3 with a median rank of 1474. This substantial failure validates the necessity of incorporating the nonlinear engine. A comparison between MSE Stack and GR-SAFS indicates that the diversity penalty improved Rank Median by 49 positions in S2 (604 → 555) while incurring a negligible AUPRC cost (< 0.01). This confirms that the Ledoit–Wolf penalty effectively enforces residual decorrelation. The full numerical details are provided in the Supplementary Material, Table S2.

**Fig. 3:**
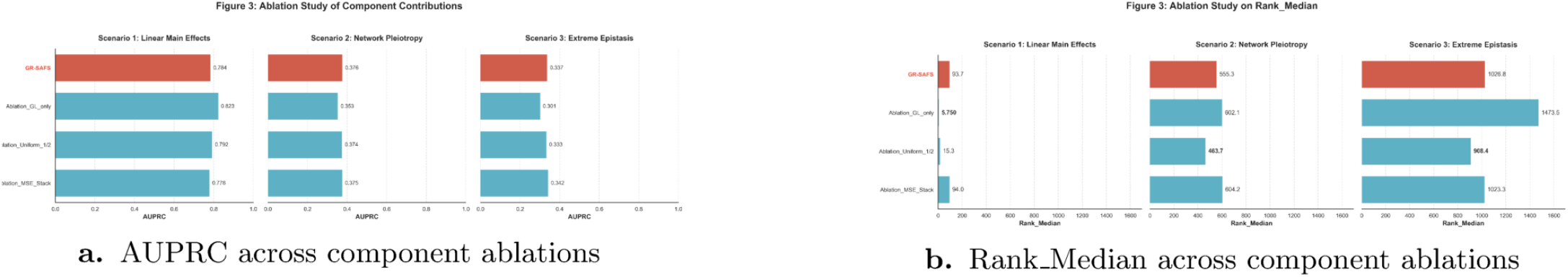
Component ablation across three scenarios. (a) AUPRC and (b) Rank Median for four variants: GL only (single-engine), Uniform (equal-weight), MSE Stack (QP with *γ* = 0), and full GR-SAFS. GR-SAFS achieves the optimal cross-scenario balance, with the diversity penalty *γ* = 10 recovering the rank stability that is otherwise lost by pure accuracy-driven weighting.

### C. GR-SAFS Identifies a 20-Gene Prognostic Signature

On the TCGA-LUAD cohort, the QP router assigned nearly balanced weights 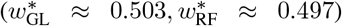, which indicates an approximately equivalent prognostic contribution from linear and nonlinear signals. The Graph-Lasso engine retained 79 non-zero genes (99.2% sparsity); following the eCDF fusion, the top 20 candidates were selected (Table II, Fig. 4). DKK1 ranked first in both engines, exhibiting strong dual-core consistency. Among the 20 candidates, 13 displayed positive coefficients (risk factors, including DKK1, FLNC, and TLE1) and 7 displayed negative coefficients (protective factors, including FAM117A and ZNF266).

**TABLE II:** Top 10 candidate prognostic genes identified by GR-SAFS on TCGA-LUAD. GL rank and RF rank denote the individual engine rankings. Direction is based on the sign of the Graph-Lasso coefficient (↑ = risk, positive; ↓ = protective, negative).

| Rank | Gene | GL rank | RF rank | Direction |
| --- | --- | --- | --- | --- |
| 1 | DKK1 | 1 | 1 | Risk ↑ |
| 2 | FLNC | 3 | 6 | Risk ↑ |
| 3 | SACK1A | 7 | 2 | Risk ↑ |
| 4 | VEGFC | 17 | 4 | Risk ↑ |
| 5 | KRT6A | 8 | 23 | Risk ↑ |
| 6 | GSEC | 11 | 20 | Risk ↑ |
| 7 | C1QTNF6 | 22 | 10 | Risk ↑ |
| 8 | FAM117A | 19 | 14 | Protect ↓ |
| 9 | ZNF266 | 30 | 8 | Protect ↓ |
| 10 | GAPDH | 43 | 7 | Risk ↑ |

**Fig. 4:**
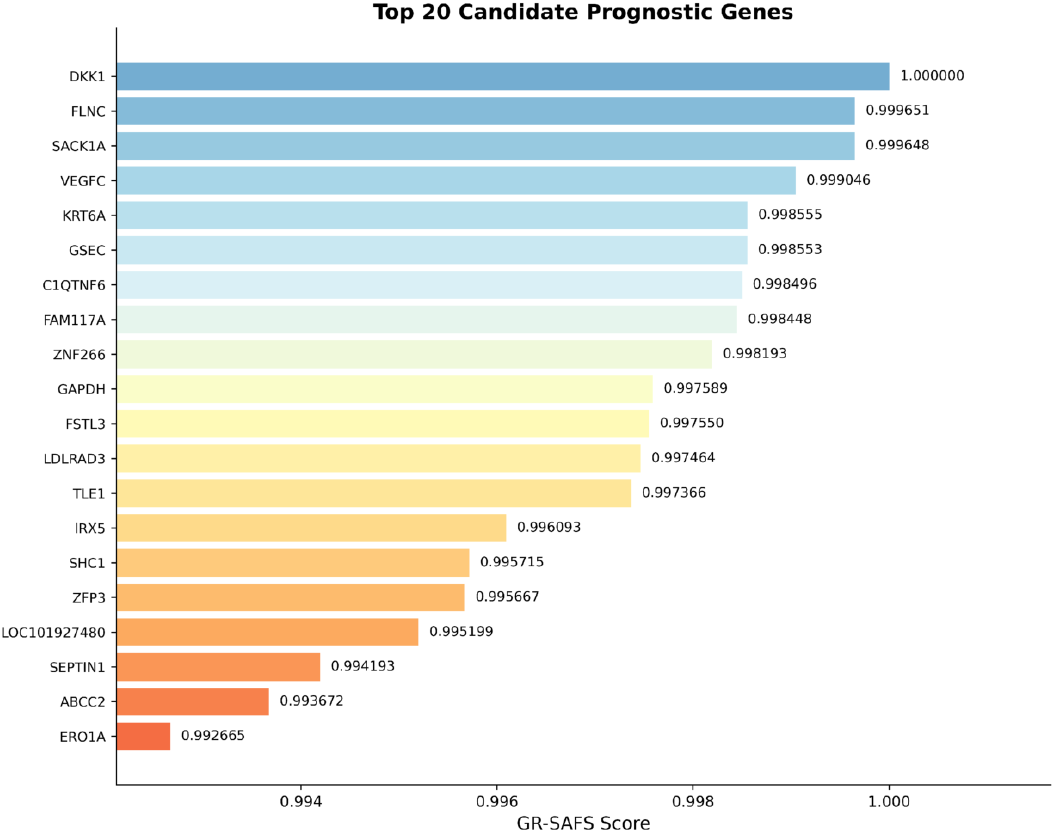
Top 20 candidate prognostic genes on TCGA-LUAD. Genes are ranked by the fused eCDF score. Red bars indicate risk factors and blue bars indicate protective factors according to the signs of the Graph-Lasso coefficients. DKK1 achieves the highest score, demonstrating strong dual-core consistency.

### D. Three-Cohort Frozen Signature Validation

Based on the direction-weighted risk score of the top 20 genes, GR-SAFS achieved highly significant stratification on TCGA-LUAD (log-rank *p* = 9.10×10^−11^, C-index = 0.700, 1-year AUC = 0.748). Under the strict frozen-signature validation, GR-SAFS maintained consistent significance across both independent cross-platform cohorts (Fig. 5, Table III): GSE31210 (*p* = 9.71× 10^−3^, C-index = 0.653) and GSE50081 (*p* = 3.68 ×10^−2^, C-index = 0.557). The Kaplan–Meier curves displayed a clear separation between the high-risk and low-risk groups from early follow-up, and ROC analysis confirmed a 1-year AUC above 0.64 across all cohorts.

**TABLE III:** Three-cohort frozen signature validation. Risk stratification remains significant (*p <* 0.05) across all cohorts.

| Cohort | $n$ | C-index | log-rank $p$ | 1y-AUC |
| --- | --- | --- | --- | --- |
| TCGA-LUAD | 504 | 0.700 | $9.10 \times 10^{-11}$ | 0.748 |
| GSE31210 | 226 | 0.653 | $9.71 \times 10^{-3}$ | 0.675 |
| GSE50081 | 181 | 0.557 | $3.68 \times 10^{-2}$ | 0.648 |

**Fig. 5:**
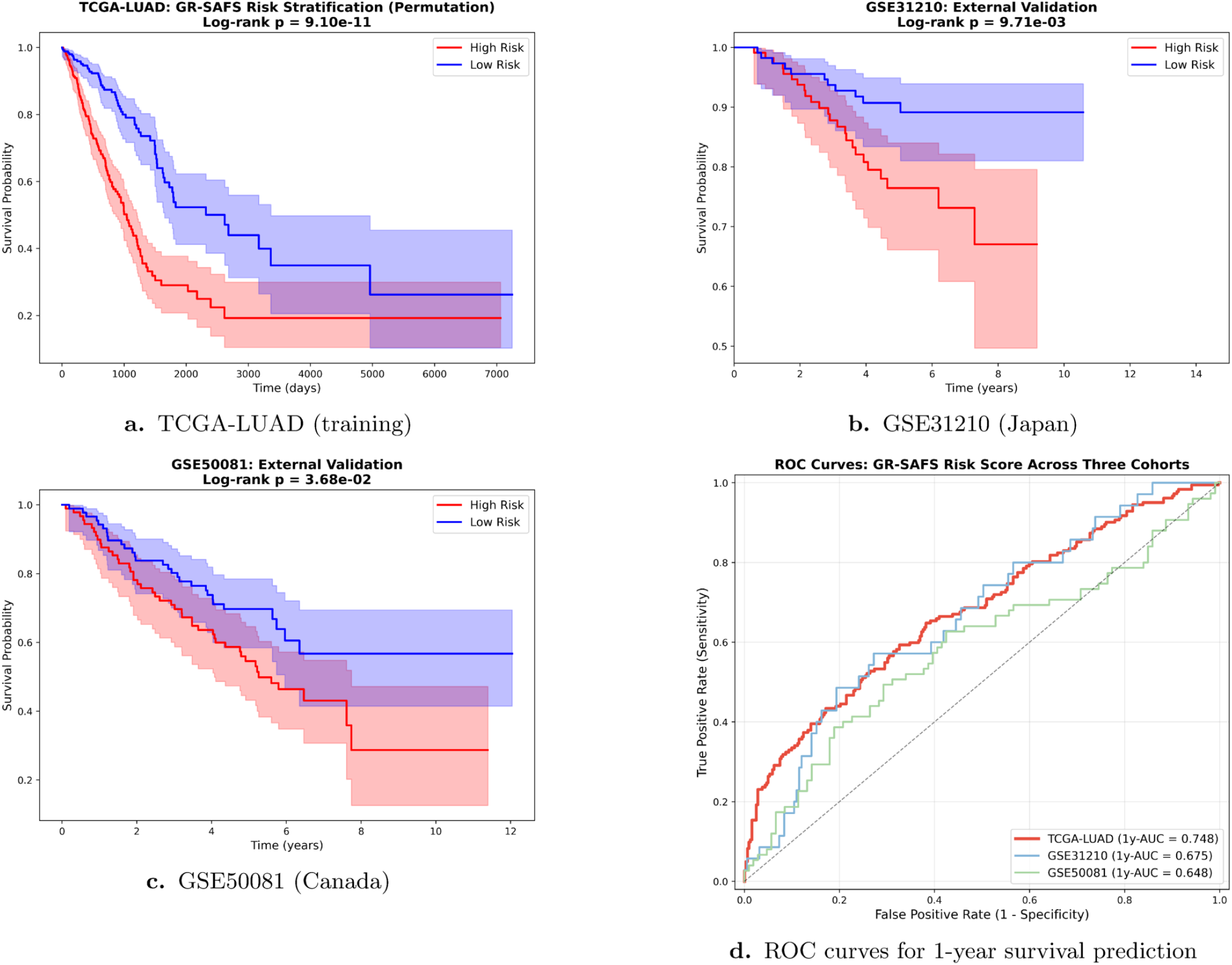
Survival analysis across three cohorts. (a)–(c) Kaplan–Meier survival curves stratified by the GR-SAFS median risk score for TCGA-LUAD (training), GSE31210 (Japan), and GSE50081 (Canada). The x-axis varies due to differences in data. (d) ROC curves for 1-year survival prediction across the three cohorts.

While GR-SAFS maintained consistent risk stratification across the cohorts, a performance decline was observed on the GSE50081 cohort (C-index = 0.557) relative to the TCGA and GSE31210 datasets. A 1000-resample bootstrap placed this C-index at 0.557 (95% confidence interval [0.48, 0.64]); the interval includes 0.5, so on this cohort the ranking metric alone is not significantly above chance, although 93% of bootstrap replicates exceeded 0.5 and the median-split risk stratification remained significant (log-rank *p* = 3.68 × 10^−2^). The GSE50081 cohort is characterized by significant clinical heterogeneity, including varied histological subtypes and a relatively small sample size. Such heterogeneity frequently introduces extreme covariate shifts when validating signatures derived from large-scale RNA-seq data like TCGA. However, despite the lower absolute performance metrics, GR-SAFS remained the only framework among the tested baselines to maintain statistical significance (*p* = 3.68 × 10^−2^) on this challenging dataset. This relative resilience suggests that the dual-core architecture and the eCDF alignment layer captured a core survival signal that persists even under high-noise conditions, prioritizing the reliability of the prognostic lower bound for cross-platform clinical deployment.

### E. Comparison with Survival Baselines

To validate GR-SAFS in high-dimensional feature selection, it was compared against seven representative models: linear shrinkage models (Cox-Lasso, Cox-EN), the univariate model (Uni-Cox), the network-constrained model (NC-Lasso), and advanced nonlinear and deep learning models (CoxBoost, DeepSurv, RSF). Under the strict frozen-signature paradigm, the results revealed significant discrepancies among architectures in handling cross-platform heterogeneity (Table IV, Fig. 6).

**TABLE IV:** Cross-platform robustness comparison. Panel (a): performance on the TCGA-LUAD training cohort. Panel (b): frozen-signature validation on two independent external cohorts. All methods employ an identical 20-gene budget. F1, Precision, and Recall are computed at the median-risk threshold. The log-rank *p* tests median-split risk stratification, all methods scored under the identical frozen-signature protocol; *p <* 0.05 denotes significant stratification. The optimal C-index/AUC/F1 value in each column is in bold; the GR-SAFS row is underlined. GR-SAFS is the only method with *p <* 0.05 in all three cohorts: Uni-Cox attains a higher GSE50081 C-index yet loses significance there (*p* = 0.14), and every other baseline fails on at least one external cohort. *n* genes: number of matched genes between the signature and each validation platform.

| Cohort | Model | C-index | $p$ (log-rank) | 1y-AUC | F1 | Precision | Recall | $n\_genes$ |
| --- | --- | --- | --- | --- | --- | --- | --- | --- |
| *(a) TCGA-LUAD training cohort* |  |  |  |  |  |  |  |  |
| | Cox-Lasso | 0.667 | $6.6 \times 10^{-9}$ | 0.654 | 0.493 | 0.425 | 0.588 | – |
| | Cox-EN | 0.686 | $1.4 \times 10^{-9}$ | 0.702 | 0.507 | 0.437 | 0.604 | – |
| | Uni-Cox | 0.697 | $9.1 \times 10^{-11}$ | 0.734 | 0.539 | 0.464 | 0.643 | – |
| | NC-Lasso | **0.712** | $1.4 \times 10^{-10}$ | **0.761** | **0.553** | **0.476** | 0.659 | – |
|  | CoxBoost | 0.589 | 0.081 | 0.650 | 0.252 | 0.154 | **0.684** | – |
|  | DeepSurv | 0.587 | 0.217 | 0.631 | 0.227 | 0.139 | 0.614 | – |
|  | RSF | 0.500 | NA | 0.500 | 0.203 | 0.113 | 1.000 | – |
| | **GR-SAFS** | <u>0.700</u> | $9.1 \times 10^{-11}$ | <u>0.748</u> | **0.553** | **0.476** | <u>0.659</u> | – |
| *(b) External validation cohorts (frozen signature)* |  |  |  |  |  |  |  |  |
| **GSE31210** | Cox-Lasso | 0.495 | 0.946 | 0.420 | 0.230 | 0.150 | 0.486 | 14 |
|  | Cox-EN | 0.537 | 0.209 | 0.557 | 0.284 | 0.186 | 0.600 | 15 |
| | Uni-Cox | **0.698** | $1.1 \times 10^{-5}$ | **0.818** | **0.392** | **0.257** | **0.829** | 20 |
|  | NC-Lasso | 0.546 | 0.540 | 0.592 | 0.257 | 0.168 | 0.543 | 12 |
|  | CoxBoost | 0.565 | 0.160 | 0.671 | 0.051 | 0.027 | <u>0.750</u> | 17 |
|  | DeepSurv | 0.595 | 0.019 | <u>0.788</u> | 0.068 | 0.035 | 1.000 | 19 |
|  | **GR-SAFS** | <u>0.653</u> | **0.010** | <u>0.675</u> | <u>0.324</u> | <u>0.212</u> | 0.686 | 19 |
| **GSE50081** | Cox-Lasso | 0.452 | 0.294 | 0.471 | 0.422 | 0.385 | 0.467 | 14 |
|  | Cox-EN | 0.475 | 0.890 | 0.484 | 0.470 | 0.429 | 0.520 | 15 |
|  | Uni-Cox | **0.567** | 0.140 | 0.647 | 0.530 | 0.484 | 0.587 | 20 |
|  | NC-Lasso | 0.501 | 0.844 | 0.569 | 0.470 | 0.429 | 0.520 | 12 |
|  | CoxBoost | 0.492 | 0.310 | NA | 0.422 | 0.385 | 0.467 | 17 |
|  | DeepSurv | 0.535 | 0.147 | NA | 0.518 | 0.473 | 0.573 | 19 |
|  | **GR-SAFS** | <u>0.557</u> | **0.037** | **0.648** | **0.554** | **0.506** | **0.613** | 19 |

**Fig. 6:**
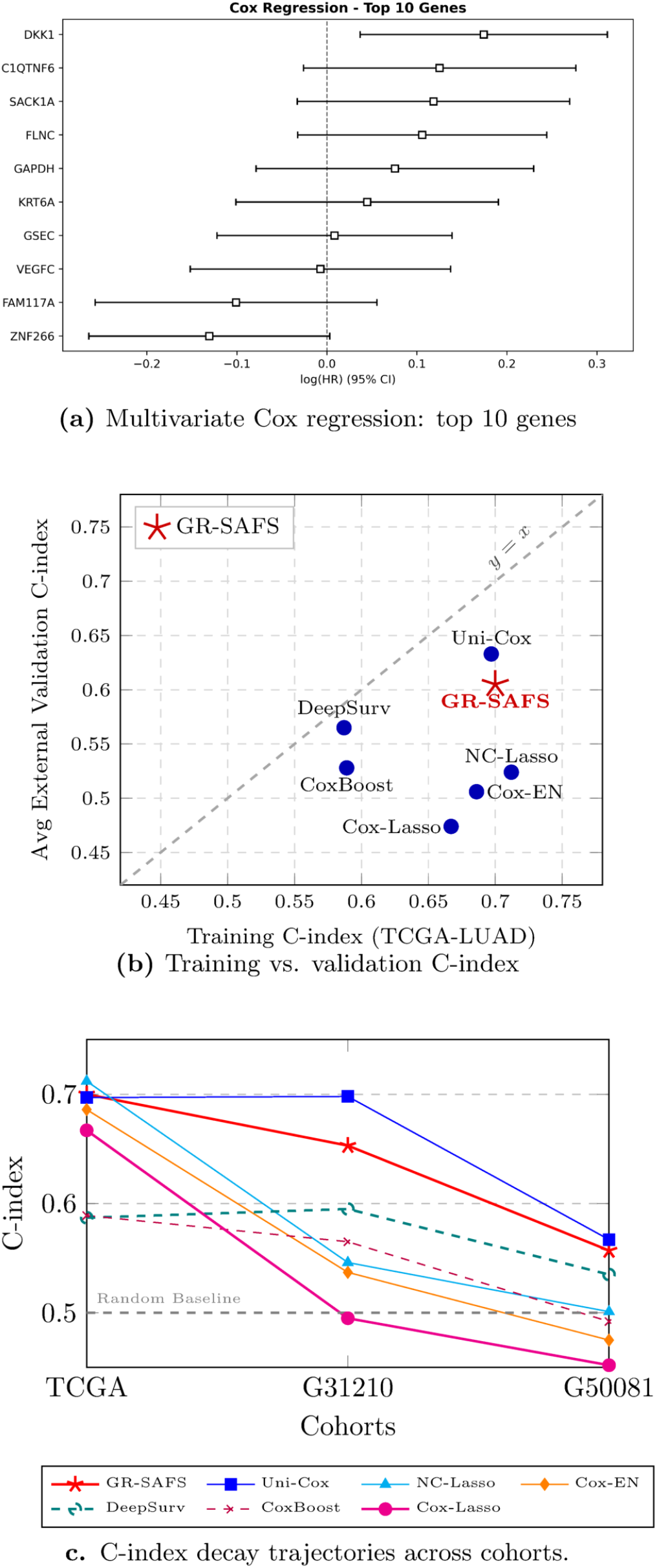
Comparison with seven survival baselines. (a) Multivariate Cox regression: top 10 genes. (b) Training vs. external validation C-index across baselines. (c) Cross-platform performance from TCGA to GSE50081, illustrating that high-C-index baselines lose significant stratification on the hardest cohort while GR-SAFS does not.

Purely data-driven nonlinear models exhibited substantial degradation in high-dimensional, small-sample regimes. Cox-Boost and DeepSurv achieved OOF C-indices of only 0.589 and 0.587, respectively, on the TCGA training set, failing to attain significant risk stratification (*p <* 0.05). The RSF model degraded when restricted to 20 features, with its OOF C-index on the TCGA training set dropping to 0.500; in its native full-feature usage RSF reaches a C-index of 0.59–0.61, so the 0.500 value reflects the 20-feature budget rather than a misconfiguration (Supplementary Material, Section S9.3).

Traditional linear models (Cox-Lasso, Cox-EN, NC-Lasso) demonstrated initial training fit but lost it under cross-platform validation, with no significant median-split stratification on either external cohort (*p >* 0.2 in all cases; Table IV).

The most informative comparison is GR-SAFS versus Uni-Cox, and we state the trade-off plainly. We do not claim that GR-SAFS attains the highest C-index: on each individual cohort at least one baseline matches or exceeds it, and Uni-Cox in particular reaches a higher external C-index (0.698 on GSE31210, 0.567 on GSE50081). What distinguishes GR-SAFS is the *reliability* of its stratification. It is the only method whose frozen signature produces statistically significant risk separation in all three cohorts (log-rank *p* = 9.1 × 10^−11^, 9.7 ×10^−3^, and 3.7 ×10^−2^). Uni-Cox, despite its higher GSE50081 C-index, loses significance on that cohort (*p* = 0.14); DeepSurv is significant on only one external cohort, and the remaining baselines on none. For a signature that is frozen and deployed on a new platform without retraining, this all-cohort significance, rather than a marginal C-index lead, is the clinically decisive property. It follows from the design: the Graph-Lasso topology prior filters platform-specific noise and the eCDF alignment limits the influence of extreme values, so the 20-gene signature transfers with a controlled performance decay instead of the collapse or loss of significance seen in the single-paradigm baselines (Fig. 6).

### F. Biological Functional Validation

To assess the biological coherence of the 20-gene signature, we performed protein–protein interaction (PPI) analysis via STRING v12.0 and functional enrichment across KEGG pathways and the three Gene Ontology (GO) categories (Biological Process, Molecular Function, Cellular Component).

### Statistical framework

Because the candidate list is small and pre-curated, single-pathway p-values are underpowered after Benjamini–Hochberg correction; we therefore report nominal p-values here, with FDR-adjusted values in the Supplementary Material (Tables S5–S7). The biological claim rests not on any single test but on the convergence of three orthogonal lines of evidence: the permutation-based PPI network-density test, the multi-ontology convergence of the Wnt/*β*-catenin signal, and independent literature support for each Top-10 gene.

### Protein–protein interaction network

The Top-20 candidates exhibited significantly more PPI edges than expected by chance (*p* = 0.003, 24 observed vs. 13 expected edges), with DKK1 and VEGFC as central hubs (Fig. 7a). The network density indicates that the selected genes participate in a coordinated interaction module rather than forming an arbitrary collection. **KEGG pathway enrichment**. KEGG enrichment (Fig. 7b) identified eight significant pathways (*p <* 0.05), the strongest being Bacterial Invasion of Epithelial Cells (*p* = 2.7 ×10^−3^), Relaxin Signaling Pathway (*p* = 6.9 × 10^−3^), and Wnt Signaling Pathway (*p* = 1.23 × 10^−2^). Although named after a microbial context, the bacterial-invasion module describes cytoskeletal remodeling, focal-adhesion dynamics, and cell-junction disassembly—machinery directly co-opted in LUAD invasion and metastasis. Consistent with this, Focal Adhesion (*p* = 1.66 ×10^−2^) and Ras Signaling Pathway (*p* = 2.20×10^−2^) were also enriched. Together these pathways indicate a coordinated invasion-and-signaling axis rather than a diffuse collection of unrelated genes.

**Fig. 7:**
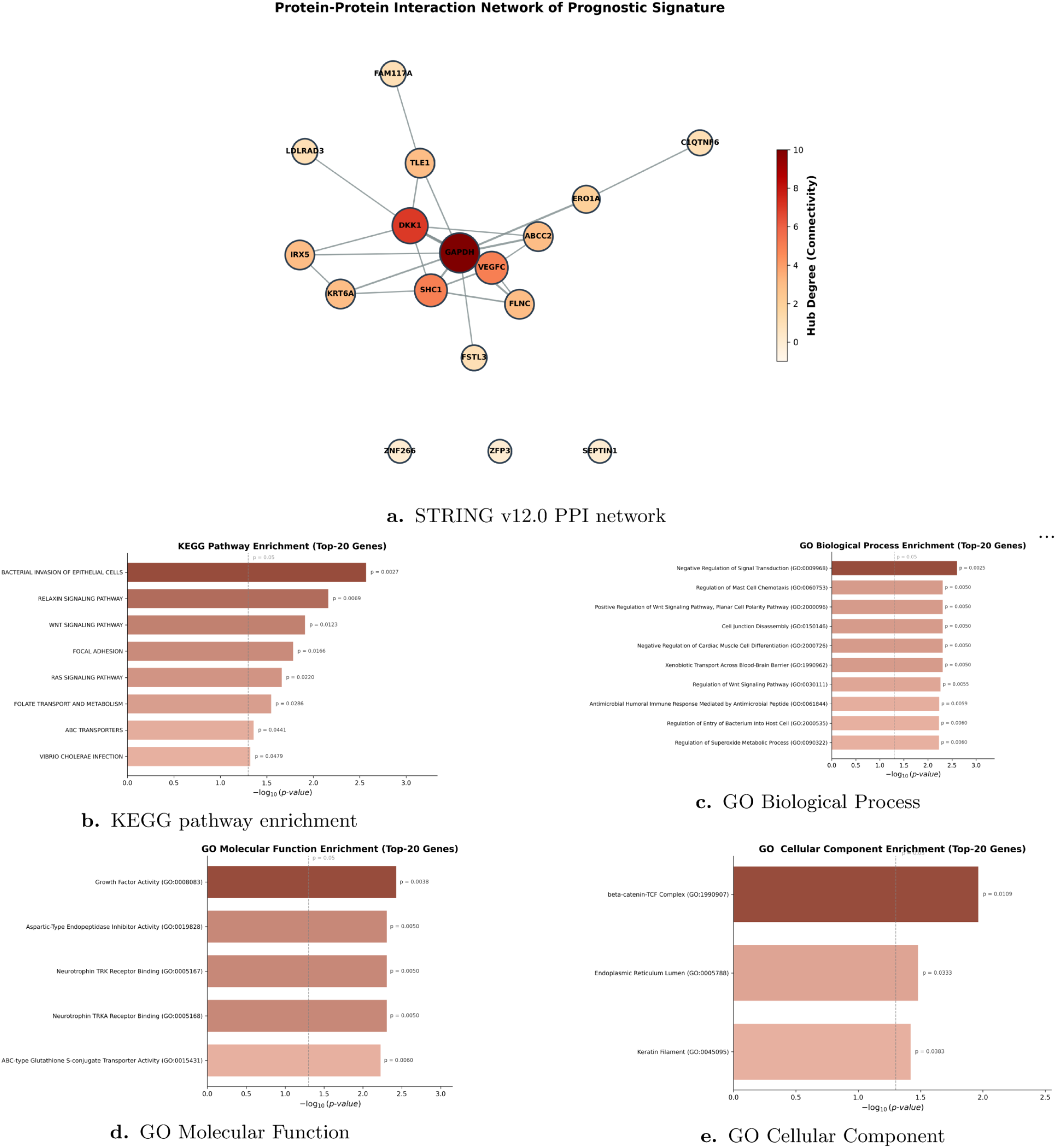
Biological functional validation of the GR-SAFS 20-gene signature. (a) STRING v12.0 PPI network. (b)–(e) Functional enrichment across KEGG pathways, GO Biological Process, GO Molecular Function, and GO Cellular Component.

### GO enrichment cross-validates a Wnt/*β*-catenin axis

The Wnt finding from KEGG was independently corroborated by GO Biological Process and GO Cellular Component (Fig. 7), a three-way multi-ontology convergence. In GO BP, the most significant term was Negative Regulation of Signal Transduction (*p* = 2.5× 10^−3^), followed by Positive Regulation of Wnt Signaling Pathway, Planar Cell Polarity Pathway (GO:2000096, *p* = 5.0 ×10^−3^) and Regulation of Wnt Signaling Pathway (GO:0030111, *p* = 5.5 ×10^−3^). In GO CC, the most significant component was the *β*-catenin-TCF Complex (GO:1990907, *p* = 1.09× 10^−2^), the canonical transcriptional effector of Wnt signaling. Because DKK1 is a secreted antagonist of Wnt/*β*-catenin, the simultaneous enrichment of the Wnt pathway, Wnt-related GO BP terms, and the *β*-catenin-TCF complex anchors the signature to a specific, mechanistically coherent signaling axis.

### GO Molecular Function

GO MF enrichment was led by Growth Factor Activity and Neurotrophin TRK Receptor Binding, consistent with the secreted ligands DKK1 and VEGFC in the signature, while Keratin Filament (driven by KRT6A) points to cytoskeletal remodeling as a complementary axis (full GO results in Supplementary Table S6).

### Literature support for the Top-10 genes

Among the Top-10 candidates, the role of well-characterized markers such as DKK1 (Wnt antagonist), VEGFC (angiogenesis), and KRT6A (cytoskeletal remodeling) in LUAD progression is well established. The signature additionally includes FLNC and SACK1A, both with prior reports in lung cancer contexts and strong dual-core consistency. The complete per-gene literature cross-reference is provided in the Supplementary Material, Section S10. The simultaneous presence of structural cytoskeletal genes (FLNC), growth-factor ligands (DKK1, VEGFC), and Wnt-signaling regulators in the signature is consistent with a coordinated invasion-signaling program rather than a single-pathway phenotype, demonstrating that the heterogeneous dual-core scanning captures functional relationships that single-engine methods would likely fragment.

## IV. Discussion

The GR-SAFS framework integrates graph-regularized priors, eCDF nonparametric alignment, and diversity-penalized QP to unify sparsity, topology awareness, and signal complementarity within high-dimensional feature selection. Strict frozen-signature validation on TCGA-LUAD coupled with two independent cross-platform cohorts confirms the generalization robustness of the framework. The nearly balanced weight allocation (*w*_GL_ ≈*w*_RF_ ≈0.5) on actual LUAD data is consistent with the hypothesis that linear main effects and nonlinear interactions contribute approximately equivalently to prognosis, motivating the dual-core architecture. Any single-paradigm method would systematically forfeit roughly half of the prognosis-relevant information, consistent with the cross-platform collapse observed in NC-Lasso and Cox-Lasso.

The simulation results also delimit where GR-SAFS is appropriate. When causal variants are independent and additive, as in scenario S1, the graph-Laplacian prior smooths coefficients across edges that carry no signal, introducing a bias that pure sparse regressors avoid. GR-SAFS is therefore intended for settings in which biological network structure is present; under a purely additive architecture, Lasso, Precision Lasso, or Knockoffs remain preferable. What the dual-core design provides is not a uniform accuracy advantage but stability as interaction and pleiotropy grow.

The limitations of this study include: the single-cancer validation on LUAD, which necessitates cross-cancer assessment; external validation restricted to Affymetrix microarray platforms; the static co-expression network construction, which lacks subtype-specific dynamic regulation; and the heuristic weight transfer from prediction space to feature space, which currently lacks strict mathematical equivalence but agrees with a model-agnostic permutation importance at the top of the ranking (Supplementary Material, Section S5.2). Future work will focus on multi-cancer generalization, dynamic networks integrated with multi-omics data, and direct Cox partial likelihood optimization.

## V. Conclusion

This study introduces GR-SAFS, a unified framework that addresses the triple challenge of sparsity, biological topology preservation, and complementary signal integration in high-dimensional prognostic biomarker discovery. Through heterogeneous dual-core scanning, eCDF distribution alignment, and diversity-penalized QP routing, GR-SAFS achieves robust cross-platform generalization on three independent LUAD cohorts. The identified 20-gene signature exhibits coherent biological interpretation through multi-ontology enrichment convergence on the Wnt/*β*-catenin axis. The framework provides a general, reproducible computational paradigm for high-dimensional feature selection in complex biomedical contexts.

## Code and Data Availability

A reproducible open-source implementation of GR-SAFS, including the preprocessing pipeline, the three engines, the eCDF/QP fusion, and scripts that regenerate all main-text figures, is available at https://github.com/hejinzhao7777-cmyk/GR-SAFS. The TCGA-LUAD data are from UCSC Xena, the validation cohorts from GEO (GSE31210, GSE50081), and clinical endpoints from TCGA-CDR; all are publicly accessible.

## Notes

### Competing Interest Statement

The authors have declared no competing interest.

## References

[1] H. Sung, J. Ferlay, R. L. Siegel et al., “Global cancer statistics 2020: GLOBOCAN estimates of incidence and mortality worldwide for 36 cancers in 185 countries,” CA: A Cancer Journal for Clinicians, vol. 71, no. 3, pp. 209–249, 2021.

[2] F. Skoulidis and J. V. Heymach, “Co-occurring genomic alterations in non-small-cell lung cancer biology and therapy,” Nature Reviews Cancer, vol. 19, no. 9, pp. 495–509, 2019.

[3] R. Tibshirani, “Regression shrinkage and selection via the lasso,” Journal of the Royal Statistical Society: Series B, vol. 58, no. 1, pp. 267–288, 1996.

[4] H. Zou and T. Hastie, “Regularization and variable selection via the elastic net,” Journal of the Royal Statistical Society: Series B, vol. 67, no. 2, pp. 301–320, 2005.

[5] L. Breiman, “Random forests,” Machine Learning, vol. 45, no. 1, pp. 5–32, 2001.

[6] H. Ishwaran, U. B. Kogalur, E. H. Blackstone, and M. S. Lauer, “Random survival forests,” The Annals of Applied Statistics, vol. 2, no. 3, pp. 841–860, 2008.

[7] S. Wiegrebe, P. Kopper, R. Sonabend et al., “Deep learning for survival analysis: A review,” Artificial Intelligence Review, vol. 57, no. 3, p. 65, 2024.

[8] Y. Deng, J. Guan, H. Guan, and T. Yang, “Multiply robust estimation for partially linear additive quantile model with missing data,” Journal of Statistical Theory and Practice, vol. 20, no. 1, p. 1, 2026.

[9] C. Li and H. Li, “Network-constrained regularization and variable selection for analysis of genomic data,” Bioinformatics, vol. 24, no. 9, pp. 1175–1182, 2008.

[10] J. Tang, M. Mou, Y. Wang et al., “dwgLASSO: An algorithm for network-based biomarker discovery,” Briefings in Bioinformatics, vol. 21, no. 6, pp. 2153–2167, 2020.

[11] F. L. Opolka, Y.-C. Zhi, P. Liò, and X. Dong, “Graph classification Gaussian processes via spectral features,” in Proceedings of the Thirty-Ninth Conference on Uncertainty in Artificial Intelligence (UAI), ser. Proceedings of Machine Learning Research, vol. 216, 2023, pp. 1575–1585.

[12] M. J. Goldman, B. Craft, M. Hastie et al., “Visualizing and interpreting cancer genomics data via the xena platform,” Nature Biotechnology, vol. 38, no. 6, pp. 675–678, 2020.

[13] H. Okayama, T. Kohno, Y. Ishii et al., “Identification of genes upregulated in ALK-positive and EGFR/KRAS/ALK-negative lung adenocarcinomas,” Cancer Research, vol. 72, no. 1, pp. 100–111, 2012.

[14] S. D. Der, J. Sykes, M. Pintilie et al., “Validation of a histology-independent prognostic gene signature for early-stage non-small-cell lung cancer,” Journal of Thoracic Oncology, vol. 9, no. 1, pp. 59–64, 2014.

[15] J. Liu, T. Lichtenberg, K. A. Hoadley et al., “An integrated TCGA pan-cancer clinical data resource to drive high-quality survival outcome analytics,” Cell, vol. 173, no. 2, pp. 400–416, 2018.

[16] T. M. Therneau, P. M. Grambsch, and T. R. Fleming, “Martingale-based residuals for survival models,” Biometrika, vol. 77, no. 1, pp. 147–160, 1990.

[17] D. H. Wolpert, “Stacked generalization,” Neural Networks, vol. 5, no. 2, pp. 241–259, 1992.

[18] O. Sagi and L. Rokach, “Ensemble learning: A survey,” Wiley Interdisciplinary Reviews: Data Mining and Knowledge Discovery, vol. 8, no. 4, p. e1249, 2018.

[19] O. Ledoit and M. Wolf, “A well-conditioned estimator for large-dimensional covariance matrices,” Journal of Multivariate Analysis, vol. 88, no. 2, pp. 365–411, 2004.

[20] S. Boyd and L. Vandenberghe, Convex Optimization. Cambridge, UK: Cambridge University Press, 2004.

[21] A. Bommert, T. Welchowski, M. Schmid, and J. Rahnenführer, “Bench-mark of filter methods for feature selection in high-dimensional gene expression survival data,” Briefings in Bioinformatics, vol. 23, no. 1, p. bbab354, 2022.

[22] M. Yuan and Y. Lin, “Model selection and estimation in regression with grouped variables,” Journal of the Royal Statistical Society: Series B, vol. 68, no. 1, pp. 49–67, 2006.

[23] N. Simon, J. Friedman, T. Hastie, and R. Tibshirani, “A sparse-group lasso,” Journal of Computational and Graphical Statistics, vol. 22, no. 2, pp. 231–245, 2013.

[24] H. Wang, B. J. Lengerich, B. Aragam, and E. P. Xing, “The precision-lasso: Accounting for correlations and linear dependencies in high-dimensional genomic data,” Bioinformatics, vol. 35, no. 7, pp. 1181–1187, 2019.

[25] R. F. Barber and E. J. Candès, “Controlling the false discovery rate via knockoffs,” The Annals of Statistics, vol. 43, no. 5, pp. 2055–2085, 2015.

[26] G. P. Way and C. S. Greene, “Extracting a biologically relevant latent space from cancer transcriptomes with variational autoencoders,” Pacific Symposium on Biocomputing, vol. 23, pp. 80–91, 2018.

[27] R. Tibshirani, “The lasso method for variable selection in the Cox model,” Statistics in Medicine, vol. 16, no. 4, pp. 385–395, 1997.

[28] T. Hothorn, P. Bühlmann, S. Dudoit, A. Molinaro, and M. J. Van Der Laan, “Survival ensembles,” Biostatistics, vol. 7, no. 3, pp. 355–373, 2006.

[29] J. L. Katzman, U. Shaham, A. Cloninger, J. Bates, T. Jiang, and Y. Kluger, “DeepSurv: Personalized treatment recommender system using a Cox proportional hazards deep neural network,” BMC Medical Research Methodology, vol. 18, no. 1, p. 24, 2018.

[30] H. Huang, J. Guan, C. Feng, J. Feng, Y. Ao, and C. Lu, “Fluid volume status detection model for patients with heart failure based on machine learning methods,” Heliyon, vol. 11, p. e41127, 2025.

