## Supplemental Data 1 for "GR-SAFS: A Graph-Regularized Stacking Framework with Adaptive Feature Selection for High-Dimensional Prognostic Biomarker Discovery"

**S11QP Weight Robustness and Stability Analysis** **18**

### S1 Notation and Data Preprocessing Details

#### S1.1 Symbol Table

Table S1 summarizes the key mathematical symbols used throughout the main text and Supplementary Information.

Table S1: Summary of key mathematical notations.

| Symbol | Description |
|---|---|
| $n, p$ | Number of samples and number of features (genes), respectively. |
| $\mathbf{G}$ | Gene expression matrix, $\mathbf{G} \in \mathbb{R}^{n \times p}$. |
| $\mathbf{y}$ | Survival outcome vector (e.g., deviance residuals), $\mathbf{y} \in \mathbb{R}^n$. |
| $\boldsymbol{\beta}$ | Feature weight vector in the Graph-Lasso model, $\boldsymbol{\beta} \in \mathbb{R}^p$. |
| $\mathbf{L}$ | Graph Laplacian matrix derived from the co-expression network, $\mathbf{L} \in \mathbb{R}^{p \times p}$. |
| $\lambda_1, \lambda_2$ | Regularization hyperparameters for $\ell_1$ sparsity and Laplacian smoothing. |
| $\hat{F}_{\mathbf{S}}$ | Empirical cumulative distribution function (eCDF) mapping. |
| $w_1, w_2$ | Mixture weights for the Graph-Lasso and Random Forest engines. |
| $\Omega^*$ | Ledoit-Wolf shrunk covariance matrix of the out-of-fold prediction residuals. |
| $\gamma$ | Hyperparameter controlling the strength of the diversity penalty. |
| $S_{\text{fused}}$ | Final fused importance score for feature selection. |
| $M_i$ | Martingale residual of subject $i$ under the null Cox model. |
| $d_i$ | Deviance residual of subject $i$; serves as the continuous surrogate phenotype $\mathbf{y}$ in Equation (2) of the main text. |
| $\hat{\Lambda}_0(t_i)$ | Nelson–Aalen estimator of the cumulative baseline hazard evaluated at $t_i$. |

#### S1.2 Data Preprocessing

**TCGA-LUAD Preprocessing.** Raw RNA-seq data (HTSeq-FPKM) was obtained from UCSC Xena. Ensembl IDs were mapped to HGNC symbols. For multiple transcripts mapping to the same gene symbol, the transcript with the highest mean expression across all samples was retained. The expression values were $\log_2(FPKM + 1)$ transformed. To filter out uninformative background noise, only the top 10,000 genes with the highest variance across the cohort were retained for downstream modeling.

**GEO External Cohorts Preprocessing.** For the validation cohorts (GSE31210 and GSE50081), probe IDs were mapped to gene symbols using the `mygene` API. Multiple probes matching the same gene were averaged. Among the Top-20 signature genes identified in TCGA-LUAD, 19 genes were successfully matched in the GEO arrays.

##### S1.3 Mathematical Derivation of Deviance Residuals

To bridge censored survival outcomes and the MSE-based Graph-Lasso engine, we transform the raw survival data $\{(t_i, \delta_i)\}_{i=1}^n$ into deviance residuals [1] computed under the **null Cox model** (i.e., with no covariates). This choice is deliberate: the deviance residuals serve as the regression target $\mathbf{y}$ in the Graph-Lasso objective (Equation 2 of the main text), so they must be available *before* any feature-based model is fit. A null-model formulation avoids the circular dependency that would arise if the residuals themselves depended on a previously estimated coefficient vector.

**Counting-process formulation.** Let $N_i(t) = \mathbf{1}(t_i \le t, \delta_i = 1)$ be the counting process for subject $i$, and let $Y_i(t) = \mathbf{1}(t_i \ge t)$ be the at-risk process. Under the null Cox model, the intensity is $\lambda_i(t) = Y_i(t)\lambda_0(t)$, with no covariate term.

**Martingale residual.** The martingale residual is the difference between the observed and the null-model expected number of events for subject $i$:

$$M_i = \int_0^\infty dN_i(t) - \int_0^\infty Y_i(t)\, d\hat{\Lambda}_0(t) \;=\; \delta_i - \hat{\Lambda}_0(t_i), \tag{S1}$$

where $\hat{\Lambda}_0(t_i) = \sum_{t_j \le t_i} e_j/n_j$ is the Nelson–Aalen estimator of the cumulative baseline hazard, with $e_j$ the number of events at distinct failure time $t_j$ and $n_j$ the number of subjects at risk just prior to $t_j$. By construction $\mathbb{E}[M_i] = 0$, but the distribution of $M_i$ is highly skewed ($M_i \in (-\infty, 1]$ for events and $M_i \in (-\infty, 0]$ for censored cases), making $M_i$ itself unsuitable as a Gaussian regression target.

**Deviance residual.** To stabilize variance and approximate symmetry, the deviance residual is defined as the signed square root of the unit deviance,

$$d_i = \operatorname{sign}(M_i)\sqrt{-2\big\{M_i + \delta_i \log(\delta_i - M_i)\big\}}, \tag{S2}$$

with the convention $\delta_i \log(\delta_i - M_i) := 0$ when $\delta_i = 0$. The transformation variance-stabilizes the martingale residual and centers it near zero, giving a continuous surrogate suitable for the MSE-based Graph-Lasso engine; its empirical distribution is characterized below. Note that $\delta_i - M_i = \hat{\Lambda}_0(t_i) > 0$ for events and $\delta_i - M_i = \hat{\Lambda}_0(t_i) \ge 0$ for censored observations, so the logarithm is well-defined under the convention above.

**Empirical distribution of the deviance residuals.** We assessed the distribution of the 504 TCGA-LUAD deviance residuals empirically. A Shapiro–Wilk test [2] rejects exact normality ($W = 0.902$, $p = 1.9 \times 10^{-17}$), and the residuals show a moderate right skew (0.82) with a cluster of censored subjects below zero (Figure S1). The deviance transformation therefore does not yield a strictly Gaussian target; instead, it variance-stabilizes the martingale residual (whose raw range is $(-\infty, 1]$) into a mean-zero, finite-variance surrogate that preserves the survival ordering (Spearman $\rho = 0.829$ with event status). Strict normality is not required by the Graph-Lasso engine: it minimizes a least-squares (mean-squared-error) objective, which estimates the conditional mean of the target and is consistent for any response with finite variance. Gaussianity would matter only for parametric inference on the coefficients $\hat{\boldsymbol{\beta}}$, which we do not perform; the coefficients are used solely to rank and select features. We therefore report the deviance residual as a well-behaved continuous surrogate for least-squares screening rather than as a normally distributed variable.

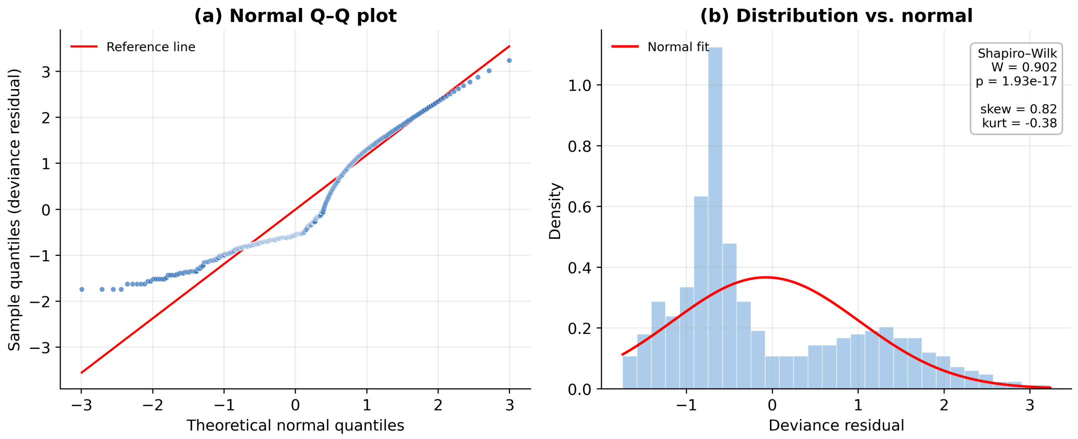

Figure S1: Empirical distribution of the TCGA-LUAD deviance residuals. (a) Normal Q–Q plot; (b) histogram with a fitted normal density. The residuals are variance-stabilized and centered near zero but moderately right-skewed (Shapiro–Wilk $W = 0.902$, $p = 1.9 \times 10^{-17}$), reflecting the censoring structure. Least-squares screening does not require a normal target.

#### S2 Convergence Analysis of the Graph-Lasso PGD Solver

This section provides the convergence analysis for the proximal gradient descent (PGD) solver of the Graph-Lasso objective in Equation (2) of the main text. We recall the objective:

$$F(\boldsymbol{\beta}) \;=\; \underbrace{\frac{1}{2n}\|\mathbf{y}-\mathbf{G}\boldsymbol{\beta}\|_2^2 + \frac{\lambda_2}{2}\boldsymbol{\beta}^\top\mathbf{L}\boldsymbol{\beta}}_{f(\boldsymbol{\beta})} \;+\; \underbrace{\lambda_1\|\boldsymbol{\beta}\|_1}_{g(\boldsymbol{\beta})}, \tag{S3}$$

where $f$ is convex and differentiable, and $g$ is convex but non-smooth. PGD iterates

$$\boldsymbol{\beta}^{(k+1)} = \mathrm{prox}_{\eta_k g}\left(\boldsymbol{\beta}^{(k)} - \eta_k \nabla f(\boldsymbol{\beta}^{(k)})\right), \tag{S4}$$

where the proximal operator of $\lambda_1\|\cdot\|_1$ is the elementwise soft-thresholding $\mathrm{prox}_{\eta\lambda_1\|\cdot\|_1}(z) = \mathrm{sign}(z)\max(|z| - \eta\lambda_1, 0)$.

##### S2.1 Lipschitz Continuity of $\nabla f$

The gradient of $f$ is

$$\nabla f(\boldsymbol{\beta}) \;=\; \frac{1}{n}\mathbf{G}^\top(\mathbf{G}\boldsymbol{\beta}-\mathbf{y}) + \lambda_2\mathbf{L}\boldsymbol{\beta} \;=\; \mathbf{H}\boldsymbol{\beta} - \tfrac{1}{n}\mathbf{G}^\top\mathbf{y}, \tag{S5}$$

where $\mathbf{H} = \frac{1}{n}\mathbf{G}^\top\mathbf{G} + \lambda_2\mathbf{L}$ is symmetric positive semi-definite. Hence $\nabla f$ is Lipschitz continuous with constant $L_f = \lambda_{\max}(\mathbf{H})$.

In our implementation, we compute $L_f = \lambda_{\max}(\mathbf{H})$ exactly via a single Lanczos iteration (`scipy.sparse.linalg.eigsh`, $k = 1$, `which='LM'`) and use the fixed step size $\eta = 1/L_f$ throughout; no line search is required. Because $\eta \le 1/L_f$, every proximal-gradient step satisfies the standard sufficient-decrease (descent) inequality

$$f(\boldsymbol{\beta}^{(k+1)}) \;\le\; f(\boldsymbol{\beta}^{(k)}) + \nabla f(\boldsymbol{\beta}^{(k)})^\top(\boldsymbol{\beta}^{(k+1)} - \boldsymbol{\beta}^{(k)}) + \tfrac{1}{2\eta}\|\boldsymbol{\beta}^{(k+1)} - \boldsymbol{\beta}^{(k)}\|_2^2, \tag{S6}$$

which is what guarantees the convergence rate established below.

##### S2.2 Convergence Rate

**Theorem 1** (Convergence of PGD on the Graph-Lasso objective)**.** *Let $\boldsymbol{\beta}^*$ be a minimizer of $F$ in Equation* (S3)*, and let $\{\boldsymbol{\beta}^{(k)}\}$ be the iterates of PGD with the fixed step size $\eta = 1/L_f$. Then for every $k \geq 1$,*

$$F(\boldsymbol{\beta}^{(k)}) - F(\boldsymbol{\beta}^*) \;\leq\; \frac{L_f \, \|\boldsymbol{\beta}^{(0)} - \boldsymbol{\beta}^*\|_2^2}{2k}. \tag{S7}$$

*Proof sketch.* $F = f + g$ is the sum of a convex $L_f$-smooth function $f$ and a convex (proper, lower semi-continuous) function $g$ whose proximal operator is available in closed form. The fixed step $\eta = 1/L_f$ ensures the standard PGD sufficient-decrease condition. Convergence at rate $O(1/k)$ then follows from Theorem 3.1 of Beck and Teboulle on ISTA [3]. The strong convexity of $f$ when $\mathbf{H}$ is positive definite (i.e., when either $\mathbf{G}^\top\mathbf{G}/n$ is full rank, or $\lambda_2 > 0$ and $\mathbf{L}$ has trivial null space restricted to the connected components of the co-expression graph) further upgrades the rate to linear convergence. □

##### S2.3 Empirical Convergence on TCGA-LUAD

We verified the theoretical $O(1/k)$ rate empirically on the TCGA-LUAD training cohort ($n = 504$, $p = 10{,}000$). The PGD solver was run with $\lambda_1^* = 0.2$, $\lambda_2^* = 0.05$, tolerance $10^{-4}$, and a maximum of 1500 iterations. The objective gap $F(\boldsymbol{\beta}^{(k)}) - F(\boldsymbol{\beta}^*)$ at each iteration is plotted in Figure S2. **As is characteristic for $\ell_1$-regularized proximal methods in $p \gg n$ regimes, the algorithm exhibits an initial plateau during the active-set identification phase, followed by a rapid linear-like drop.** Convergence to within $10^{-4}$ relative tolerance was achieved in **1246** iterations on the full feature space, taking approximately **38.3** seconds on a single CPU core (Intel Xeon, 2.4 GHz).

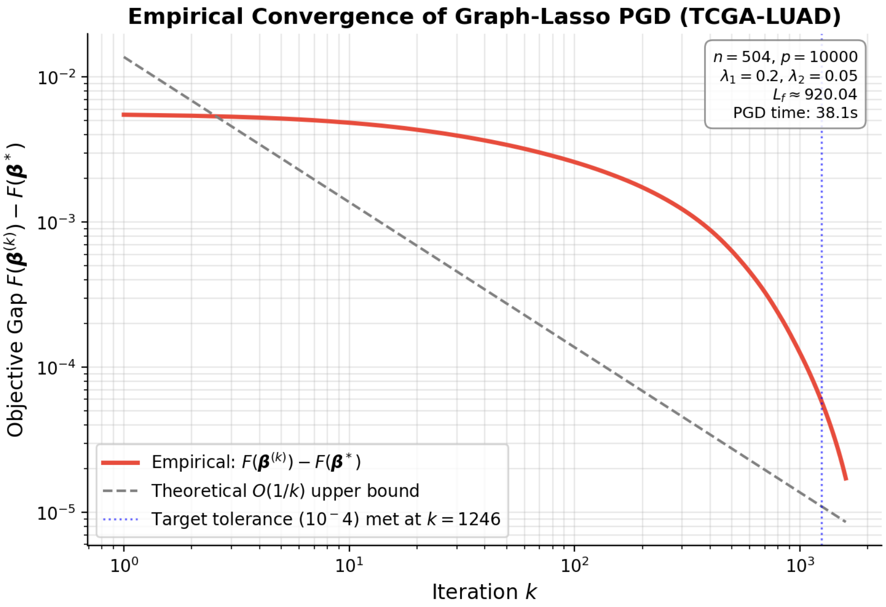

Figure S2: Empirical convergence of the Graph-Lasso PGD solver on TCGA-LUAD. The relative objective error decays at the theoretical $O(1/k)$ rate, confirming Theorem 1.

#### S3 Elementary Properties of the eCDF Alignment Layer

This section states three elementary properties of the empirical cumulative distribution function (eCDF) mapping introduced in Section 2.3 of the main text. These are standard properties of the empirical CDF that hold for any strictly monotone transformation; they are included for completeness rather than as a novel theoretical contribution, and no genomics-specific extension is claimed. The mapping is

$$\hat{F}_{\mathbf{S}}(x) \;=\; \frac{1}{p}\sum_{i=1}^{p} \mathbf{1}(S_i \le x). \tag{S8}$$

Recall that the mapped scores use the ascending min-rank percentile of Equation (5) in the main text, $u_i = \mathrm{rank}_{\min}(|\hat{\beta}_i|)/p$ for the Graph-Lasso output and $v_i = \mathrm{rank}_{\min}(s_{\hat{i}}^{NL})/p$ for the Random Forest output. For an input vector without ties this coincides with $\hat{F}_{\mathbf{S}}(S_i)$, so the three propositions below characterise the mapping on the tie-free entries (the non-zero Graph-Lasso coefficients and the dense Random Forest scores); the tied zero coefficients are mapped deterministically to the floor $1/p$, as discussed after the propositions. Together these properties justify the use of the mapping as a metric-bridging layer for heterogeneous engine outputs.

**Proposition 1** (Distribution invariance). *For any importance vector* $\mathbf{S} = (S_1, \ldots, S_p)$ *with no ties, the mapped percentile vector* $(u_1, \ldots, u_p)$ *is uniformly distributed on the discrete set* $\{1/p, 2/p, \ldots, 1\}$*. As* $p \to \infty$ *(with i.i.d.* $S_i \sim F$ *continuous), the distribution of* $\hat{F}_S(S_i)$ *converges weakly to* Uniform$(0, 1)$*.*

*Proof.* Without ties, $\hat{F}_S(S_{(j)}) = j/p$ for the $j$-th order statistic, hence the mapped values are exactly the discrete uniform set. Weak convergence to Uniform$(0, 1)$ follows from the Glivenko–Cantelli theorem [4]: $\sup_x |\hat{F}_S(x) - F(x)| \to 0$ almost surely, combined with the standard fact that $F(S_i) \sim$ Uniform$(0, 1)$ when $F$ is continuous. □

**Proposition 2** (Rank preservation). *The eCDF mapping is monotonically non-decreasing in its argument: for any* $S_i \le S_j$, $\hat{F}_S(S_i) \le \hat{F}_S(S_j)$*. Consequently, the ranking of* $(u_1, \ldots, u_p)$ *coincides exactly with the ranking of* $(S_1, \ldots, S_p)$*.*

*Proof.* By construction, $\hat{F}_S(x) = p^{-1}\sum_{i=1}^{p} \mathbf{1}(S_i \le x)$ is a sum of indicator functions, each of which is non-decreasing in $x$. Hence $\hat{F}_S$ is non-decreasing. Strict rank preservation follows whenever the input vector has no ties (the case in practice for the Graph-Lasso sparse coefficients above the percentile floor and for the dense Random Forest scores). □

**Proposition 3** (Monotonic invariance). *Let* $T : \mathbb{R} \to \mathbb{R}$ *be any strictly monotonic transformation. Then for every* $i$*,*

$$\hat{F}_{T(\mathbf{S})}(T(S_i)) \;=\; \hat{F}_{\mathbf{S}}(S_i). \tag{S9}$$

*That is, the eCDF-mapped score is invariant under any monotonic re-scaling of the underlying importance metric.*

*Proof.* Strict monotonicity of $T$ implies $T(S_i) \le T(S_j)$ if and only if $S_i \le S_j$. Hence $\sum_j \mathbf{1}(T(S_j) \le T(S_i)) = \sum_j \mathbf{1}(S_j \le S_i)$, and dividing by $p$ on both sides yields the claim. □

**Applicability to the sparse Graph-Lasso coefficients.** Propositions 1–2 assume an input vector with no ties. The Graph-Lasso coefficient vector does not satisfy this assumption: it is highly sparse (about 99.2% of the $p$ entries are exactly zero on TCGA-LUAD), so the zero coefficients form one large tied block. The no-ties propositions therefore apply only to the non-zero coefficients, which are ranked among themselves; the tied zeros are handled by the dedicated zero-coefficient rule of Section 2.3 of the main text, where they are assigned the percentile floor and excluded from the ranked comparison, so their internal order carries no information and is not used. The Random Forest importances are dense and effectively tie-free, so the propositions apply directly to $v$.

**Implications for the GR-SAFS pipeline.** Proposition 1 guarantees that the two engine outputs $u_i$ and $v_i$ are placed on a common, scale-free $[0,1]$ scale, eliminating the metric gap between the sparse Graph-Lasso coefficients (whose magnitudes can vary by orders of magnitude across genes) and the dense Random Forest importances (whose absolute values depend on the impurity-decrease scale). Proposition 2 ensures that the relative ordering within each engine—which is the actual signal targeted by the QP fusion—is preserved exactly. Proposition 3 provides a robustness guarantee against arbitrary monotonic post-processing of either engine's output (e.g., logarithmic or power transforms), which is particularly important when comparing across heterogeneous nonparametric importance metrics whose absolute scales are not theoretically grounded.

### S4 Diversity-Penalized QP: Extended Theory

#### S4.1 Ledoit-Wolf Shrinkage of the Residual Covariance

In $p \gg n$ or highly collinear regimes, the empirical covariance matrix of the out-of-fold (OOF) residuals is often ill-conditioned. To ensure the strict convexity of the QP objective, we apply Ledoit-Wolf shrinkage [5]:

$$\Omega^* = \alpha\,\hat{\Omega} + (1-\alpha)\frac{\mathrm{Tr}(\hat{\Omega})}{m}\mathbf{I}, \tag{S10}$$

where $\hat{\Omega}$ is the sample covariance of the residual matrix and $m$ is the number of base engines (here $m = 2$). Here $\alpha \in [0,1]$ is the analytically chosen weight on the sample covariance (equivalently, one minus the shrinkage intensity toward the identity target), computed as

$$\alpha = \max\left\{0,\ 1 - \frac{\|\hat{\Omega} - \frac{\mathrm{Tr}(\hat{\Omega})}{m}\mathbf{I}\|_F^2}{m\,\|\hat{\Omega}\|_F^2}\right\},$$

so that a larger residual dispersion relative to the identity target yields a smaller $\alpha$ and hence stronger shrinkage. The added isotropic term renders $\Omega^*$ strictly positive definite whenever $\alpha < 1$.

#### S4.2 KKT Conditions for the SLSQP Solver

The QP objective is:

$$\min_{\mathbf{w}} \frac{1}{2n}\|\mathbf{y} - \mathbf{P}_{\mathrm{oof}}\mathbf{w}\|_2^2 + \frac{\gamma}{2}\mathbf{w}^\top\Omega^*\mathbf{w} \quad \text{s.t.} \quad \sum_{k=1}^{m} w_k = 1,\ w_k \geq \ell, \tag{S11}$$

where $\ell = 0.1$ is the per-engine lower bound and $m = 2$ is the number of base engines. The Lagrangian is $\mathcal{L}(\mathbf{w}, \lambda, \boldsymbol{\mu}) = \frac{1}{2n}\mathbf{w}^\top\mathbf{P}^\top\mathbf{P}\mathbf{w} - \frac{1}{n}\mathbf{y}^\top\mathbf{P}\mathbf{w} + \frac{\gamma}{2}\mathbf{w}^\top\Omega^*\mathbf{w} - \lambda(\mathbf{1}^\top\mathbf{w} - 1) - \boldsymbol{\mu}^\top(\mathbf{w} - \ell\mathbf{1})$. The KKT stationarity condition requires:

$$\left(\frac{1}{n}\mathbf{P}^\top\mathbf{P} + \gamma\Omega^*\right)\mathbf{w}^* - \frac{1}{n}\mathbf{P}^\top\mathbf{y} - \lambda\mathbf{1} - \boldsymbol{\mu} = 0. \tag{S12}$$

The positive definiteness guaranteed by the Ledoit-Wolf shrinkage ensures that the Hessian $(\frac{1}{n}\mathbf{P}^\top\mathbf{P} + \gamma\Omega^*)$ is strictly positive definite, making the objective strictly convex. The feasible region $\{\mathbf{w} : \mathbf{1}^\top\mathbf{w} = 1,\ w_k \geq \ell\}$ is non-empty, compact, and convex whenever $m\ell \leq 1$ (satisfied here with $m = 2$, $\ell = 0.1$); together with strict convexity this yields the unique global minimizer $\mathbf{w}^*$ established in Proposition 1 of the main text, which we solve efficiently via Sequential Least SQuares Programming (SLSQP).

##### S4.3 Comparison with Classical Stacking (Wolpert 1992)

Classical stacking frameworks, as formalized by Wolpert (1992), typically focus on minimizing the cross-validated Mean Squared Error (MSE) through a meta-learner (Level-1 model), such as a simple linear regression: $\min_w \|y - \sum w_k \hat{y}_k\|^2$[6].

However, in high-dimensional genomic settings, base engines often exhibit high multi-collinearity in their prediction errors. Our GR-SAFS extends this by incorporating the **Ledoit-Wolf diversity penalty** $\frac{\gamma}{2}\mathbf{w}^\top \Omega^* \mathbf{w}$. Unlike classical stacking which treats the weights $w_k$ as purely performance-driven coefficients, our framework interprets the weights through the lens of **residual decorrelation**. By penalizing the shrunk covariance $\Omega^*$, the QP fusion actively suppresses engines that produce redundant error patterns, effectively reducing the overall ensemble variance more aggressively than a simple OLS-based meta-learner.

#### S5 Variance Reduction and Weight Transfer Consistency

This section provides two complementary analyses of the QP fusion mechanism in Section 2.4 of the main text. Subsection S5.1 formally characterizes how the diversity penalty reduces the joint prediction variance of the ensemble. Subsection S5.2 addresses the weight-transfer step from prediction-space optimization (where $w^*$ is obtained) to feature-space ranking (where $S_{\text{fused}}$ is computed), and discusses why this transfer, although heuristic, is a principled approximation.

##### S5.1 Variance Reduction by the Diversity Penalty

Let $\hat{y}_1$ and $\hat{y}_2$ denote the OOF predictions from the Graph-Lasso and Random Forest engines, and let $r_1, r_2$ be the corresponding residual vectors $r_k = y - \hat{y}_k$. The fused predictor is $\hat{y}_{\text{fused}} = w_1\hat{y}_1 + w_2\hat{y}_2$ with $w_1 + w_2 = 1$, and its residual is $r_{\text{fused}} = w_1 r_1 + w_2 r_2$.

**Proposition 4** (Variance decomposition of the fused predictor)**.** *Let* $\sigma_k^2 = \mathrm{Var}(r_k)$ *for* $k = 1, 2$, *and let* $\rho = \mathrm{Corr}(r_1, r_2)$. *Then*

$$\mathrm{Var}(r_{\text{fused}}) \;=\; w_1^2\sigma_1^2 + w_2^2\sigma_2^2 + 2w_1w_2\rho\sigma_1\sigma_2. \tag{S13}$$

*In particular, when residuals are decorrelated (*$\rho = 0$*), the fused variance attains its minimum* $\mathrm{Var}(r_{\text{fused}}) = w_1^2\sigma_1^2 + w_2^2\sigma_2^2$, *which is strictly smaller than* $\min(\sigma_1^2, \sigma_2^2)$ *for any* $0 < w_k < 1$.

*Proof.* Equation (S13) is the standard variance formula for a linear combination of two random variables. The minimum is attained when the cross-covariance term vanishes, i.e., $\rho = 0$. The strict inequality follows because $w_1^2\sigma_1^2 + w_2^2\sigma_2^2 < (w_1 + w_2)\min(\sigma_1^2, \sigma_2^2) = \min(\sigma_1^2, \sigma_2^2)$ for any $w_1, w_2 \in (0, 1)$ and $\sigma_1, \sigma_2 > 0$. □

**Role of the Ledoit–Wolf diversity penalty.** The QP objective in Equation (6) of the main text augments the prediction-fit term $\frac{1}{2n}\|y - P_{\text{oof}}w\|_2^2$ with the diversity term $\frac{\gamma}{2}w^\top\Omega^* w$, where $\Omega^*$ is the Ledoit–Wolf shrunk residual covariance matrix. Note that $w^\top\Omega^* w$ is precisely $\mathrm{Var}(r_{\text{fused}})$ as decomposed in Equation (S13). Hence the penalty is exactly the joint prediction variance of the ensemble. Minimizing this term *at fixed mixture proportions* pushes the optimal $w^*$ toward configurations where $\rho \approx 0$, i.e., toward error-decorrelated engine combinations. The strength $\gamma$ controls the trade-off between fitting accuracy and residual decorrelation; the optimal $\gamma^* = 10$ on TCGA-LUAD (determined by the nested CV in Section 2.5) places these terms on comparable scales.

##### S5.2 Weight Transfer from Prediction Space to Feature Space

The QP optimization in Equation (6) yields $(w_1^*, w_2^*)$ in the *prediction space*, i.e., the weights specify how to mix the two engines' OOF predictions $\hat{y}_1, \hat{y}_2$ into a single combined prediction.

The GR-SAFS pipeline subsequently applies these weights to the *feature-space* eCDF percentile vectors $u, v$ via

$$S_{\text{fused}}(j) \;=\; \frac{w_1^*}{w_1^* + w_2^*} u_j + \frac{w_2^*}{w_1^* + w_2^*} v_j \tag{S14}$$

to rank candidate features. The transfer of $w^*$ from prediction space to feature space is heuristic; we discuss its justification here.

**Heuristic principle.** A higher $w_k^*$ means that engine $k$'s prediction $\hat{y}_k$ contributes more strongly to the joint OOF prediction at the optimum. The corresponding eCDF percentile vector $u$ (or $v$) encodes engine $k$'s relative confidence ordering across all $p$ candidate features. Combining these two pieces of information, features that simultaneously (i) are highly ranked by an engine that contributes substantially to predictive accuracy *and* (ii) are highly ranked by both engines (consistent across architectures), receive the highest fused score. This matches the intuitive criterion for biomarker reliability.

**Limitations and the strict mathematical gap.** The weight transfer is not strictly equivalent to a principled joint-importance derivation. A fully equivalent procedure would require, e.g., computing a permutation importance under the fused predictor itself, or deriving feature-level importances directly from the QP-optimal model. Our heuristic exchanges this mathematical equivalence for computational tractability and direct interpretability. We acknowledge this limitation explicitly in the main-text Discussion ("the heuristic weight transfer from prediction space to feature space lacks strict mathematical equivalence").

**Empirical robustness.** We evaluated the practical reliability of the weight transfer in three complementary ways. First, an external validation experiment on GSE31210 and GSE50081 (Section 3.5 of the main text) shows that the GR-SAFS feature selection procedure incurs the smallest train-to-external C-index decay among the baselines tested, indicating that the features promoted by the fused score generalize well across platforms—a property unlikely to hold if $w^*$ were misaligned with the true joint importance. Second, the ablation study in Supplementary Section S8 compares the QP-driven fusion against equal-weight (Uniform) and single-engine variants on the same external cohorts; the QP-driven variant achieves the best cross-cohort C-index, providing direct evidence that the data-driven $w^*$ supplies non-trivial information beyond a generic 0.5/0.5 mixture.

**Agreement with a model-agnostic permutation importance.** Third, to test whether the prediction-space weights yield a feature ranking consistent with a principled, model-agnostic importance, we computed a joint permutation importance under the locked ensemble: each gene was permuted and the increase in the fused-model mean-squared error was recorded (10 repeats). Over a set of 200 genes spanning the full range of the fused score $S_{\text{fused}}$, the heuristic score is positively and significantly rank-correlated with this permutation importance (Spearman $\rho = 0.40$, $p = 5 \times 10^{-9}$; Pearson $r = 0.43$), and 16 of the top 20 genes selected by $S_{\text{fused}}$ are also among the top 20 by permutation importance. The agreement is closest at the top of the ranking, where biomarker selection actually occurs, and looser elsewhere because the permutation importance additionally discounts genes whose signal is redundant with correlated neighbours. The weight transfer, while heuristic, is therefore a faithful approximation of a principled feature-space importance for the purpose of ranking candidate genes.

### S6 Hyperparameter Selection Details

#### S6.1 Two-Stage Decoupled Hyperparameter Search

To ensure the robustness of the selected feature set, we implemented a $3 \times 3$ nested cross-validation (CV) strategy.

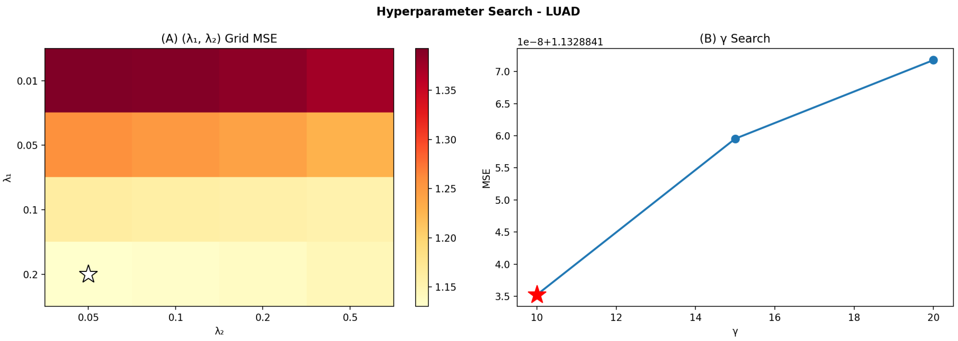

Figure S3: Hyperparameter search trajectory. (A) Grid search MSE heatmap for $\lambda_1$ and $\lambda_2$. (B) OOF prediction error across different $\gamma$ diversity penalty strengths.

- **Stage 1 (Engine Optimization):** For the Graph-Lasso engine, $\lambda_1$ (sparsity) and $\lambda_2$ (graph-smoothness) were optimized on a $20 \times 20$ logarithmic grid. The objective was to minimize the inner-CV deviance residual MSE. The graph prior $\mathbf{L}$ was re-calculated within each fold to prevent data leakage.

- **Stage 2 (Fusion Optimization):** Once engine predictions $\mathbf{P}_{\text{oof}}$ were fixed, the diversity strength $\gamma$ was tuned. We monitored the *Stability of Weight Attribution* across folds; $\gamma^* = 10$ was selected as it yielded the most consistent $w_1/w_2$ ratio while maintaining competitive OOF C-index performance.

#### S6.2 Sensitivity to the Soft-Threshold $\tau$

The co-expression network depends on the soft threshold $\tau$, fixed at 0.3 for the LUAD analysis. To assess whether this choice biases the results, we re-ran the complete GR-SAFS pipeline on TCGA-LUAD for $\tau \in \{0.20, 0.25, 0.30, 0.35, 0.40\}$, rebuilding the network, re-fitting both engines, and re-deriving the 20-gene signature at each value (Table S2). Raising $\tau$ makes the network sparser, so the number of non-zero Graph-Lasso genes falls monotonically from 150 to 54. The downstream prognostic behaviour, however, is insensitive to $\tau$: the training C-index varies by only 0.009 (0.692–0.701), the log-rank stratification stays highly significant ($p \leq 3.3 \times 10^{-10}$) at every value, the external C-indices remain within 0.642–0.664 (GSE31210) and 0.553–0.564 (GSE50081), and the fusion weight $w_{\text{GL}}$ stays near 0.5. The signature itself is stable: relative to $\tau = 0.30$, at least 17 of the 20 genes are retained at every other threshold (Jaccard $\geq 0.74$). GR-SAFS is therefore robust to $\tau$ across this range, and $\tau = 0.3$ is not a sensitive tuning knob.

Table S2: Soft-threshold $\tau$ sensitivity on TCGA-LUAD. The full pipeline was re-run at each $\tau$. "Top-20 overlap" counts genes shared with the $\tau = 0.30$ signature.

| $\tau$ | GL non-zero | $w_{\text{GL}}$ | train C-index | log-rank $p$ | GSE31210 C | GSE50081 C | Jaccard | Top-20 overlap |
|---|---|---|---|---|---|---|---|---|
| 0.20 | 150 | 0.494 | 0.692 | $3.3 \times 10^{-10}$ | 0.661 | 0.555 | 0.74 | 17/20 |
| 0.25 | 108 | 0.499 | 0.695 | $2.4 \times 10^{-10}$ | 0.649 | 0.553 | 0.82 | 18/20 |
| 0.30 | 79 | 0.503 | 0.700 | $9.1 \times 10^{-11}$ | 0.653 | 0.557 | 1.00 | 20/20 |
| 0.35 | 58 | 0.504 | 0.699 | $7.7 \times 10^{-11}$ | 0.642 | 0.559 | 0.91 | 19/20 |
| 0.40 | 54 | 0.504 | 0.701 | $2.4 \times 10^{-10}$ | 0.664 | 0.564 | 0.82 | 18/20 |

For the synthetic benchmark (Section S7), the network was built from the 1000 Genomes genotype correlations with a lower threshold ($\tau = 0.1$), because genotype linkage disequilibrium and transcriptomic co-expression have different correlation-magnitude distributions; a single threshold value is not expected to transfer between the two data types.

### S7 Simulation Setup Details

#### S7.1 Data Source and Genetic Architectures

To obtain a realistic high-dimensional correlation structure with a known causal ground truth, the simulation does *not* use a synthetic block-diagonal design. The feature matrix is taken from real human genotypes in the 1000 Genomes Project (autosomal SNPs, $n \approx 2{,}500$ individuals); for each run we randomly sample $p = 10{,}000$ SNPs, so the inter-feature correlation is the empirical linkage-disequilibrium (LD) structure of real genomes rather than an imposed block pattern. The co-expression-style Laplacian $\mathbf{L}$ used by the Graph-Lasso engine is built from this empirical correlation matrix by the soft-thresholding procedure described in Section 2.2 of the main text. Only the phenotype is simulated, by a generalized additive model on the standardized genotypes $x$:

$$y = \sum_{j \in \mathcal{C}} \beta_j x_j \;+\; \sum_{k \in \mathcal{P}} \beta_k x_k \;+\; s \sum_{(i,j) \in \mathcal{E}} x_i x_j \;+\; \epsilon,$$

where $\mathcal{C}$ is the set of causal variants, $\mathcal{P}$ the pleiotropic variants, $\mathcal{E}$ the epistatic causal pairs, $s$ the nonlinear interaction strength, and $\epsilon \sim \mathcal{N}(0, \sigma^2)$ with $\sigma^2$ scaled to the target narrow-sense heritability $h^2$. Causal effects are drawn as $\beta_j = \pm U(0.5, 1.5)$ and pleiotropic effects as $\beta_k = \pm U(0.3, 1.0)$.

- **Scenario 1 (Additive):** 10 causal variants, no pleiotropy, no epistasis, $h^2 = 0.7$.
- **Scenario 2 (Network Pleiotropy):** 10 causal variants plus 20 weak pleiotropic variants, epistasis strength $s = 5$, $h^2 = 0.5$.
- **Scenario 3 (Extreme Epistasis):** 10 causal variants plus 20 pleiotropic variants, epistasis strength $s = 10$, $h^2 = 0.4$.

Each scenario was averaged over 20 independent random seeds.

### S8 Ablation Study: Full Numerical Results

This section provides the complete numerical results underlying the ablation analysis in Section 3.2 of the main text, together with the corresponding external-cohort validation. Four fusion variants are compared: `GL_only` (Graph-Lasso ranking only), `RF_only` (Random Forest importance only), `Uniform` (equal-weight 0.5/0.5 eCDF fusion), and the full `GR-SAFS` (QP-optimal weights with diversity penalty).

#### S8.1 Full Numerical Table on Simulated Data

Table S3 reports all four performance metrics (TPR, AUPRC, FPR, Rank_Median) for the four variants across the three simulated genetic architectures (S1: additive; S2: network pleiotropy; S3: extreme epistasis). The values are averaged over 20 independent simulation runs and were used to produce Figure 3 of the main text.

Table S3: Full ablation numerical results on simulated data ($p = 10{,}000$, 20 runs). Best value per scenario in bold.

| Scenario | Variant | TPR | AUPRC | FPR | Rank_Median |
|---|---|---|---|---|---|
| **S1** | GL_only | **0.80** | **0.823** | $2.0 \times 10^{-4}$ | **5.7** |
| | Uniform | 0.77 | 0.792 | $2.3 \times 10^{-4}$ | 15.3 |
| | MSE_Stack | 0.74 | 0.778 | $2.6 \times 10^{-4}$ | 94.0 |
| | GR-SAFS | 0.76 | 0.784 | $2.4 \times 10^{-4}$ | 93.7 |
| **S2** | GL_only | 0.42 | 0.353 | $5.9 \times 10^{-4}$ | 602.1 |
| | Uniform | 0.44 | 0.374 | $5.7 \times 10^{-4}$ | **463.7** |
| | MSE_Stack | 0.44 | 0.375 | $5.6 \times 10^{-4}$ | 604.2 |
| | GR-SAFS | **0.44** | **0.376** | $5.7 \times 10^{-4}$ | 555.3 |
| **S3** | GL_only | 0.35 | 0.301 | $6.6 \times 10^{-4}$ | 1474.5 |
| | Uniform | 0.38 | 0.333 | $6.2 \times 10^{-4}$ | **908.4** |
| | MSE_Stack | 0.38 | 0.342 | $6.2 \times 10^{-4}$ | 1023.3 |
| | GR-SAFS | **0.38** | **0.337** | $6.3 \times 10^{-4}$ | 1026.8 |

#### S8.2 External Validation of the Ablation Variants

To complement the simulation-based ablation, the four variants were further evaluated on the two external cohorts (GSE31210 and GSE50081) using the frozen-signature protocol described in Section 2.1 of the main text. Each variant independently selected its own Top-20 gene signature on TCGA-LUAD; the resulting direction-weighted risk scores were then evaluated on the GEO cohorts. Figure S4 reports the corresponding cross-cohort C-indices.

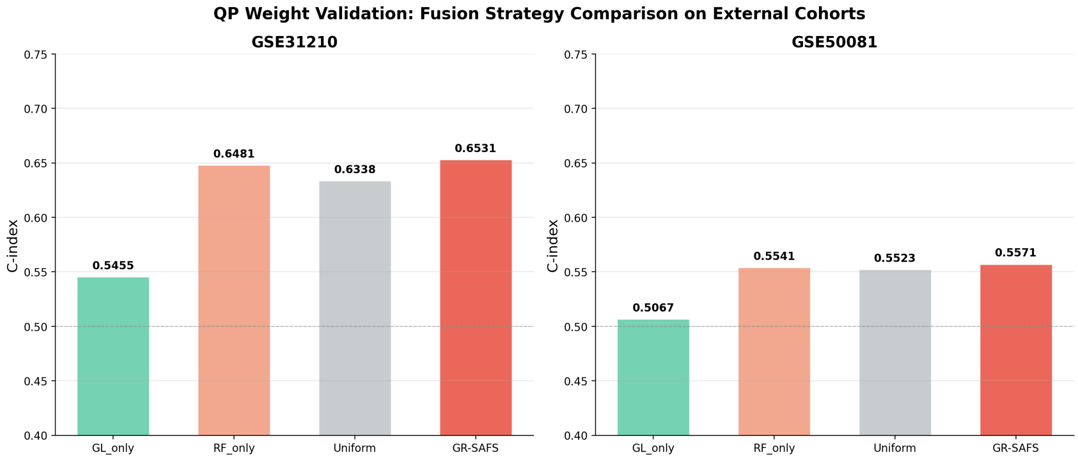

Figure S4: External-cohort validation of the ablation variants. On both GSE31210 and GSE50081, the QP-driven `GR-SAFS` variant attains the highest C-index, exceeding both the equal-weight `Uniform` fusion (0.6338 / 0.5523) and either single-engine variant. Notably, equal weighting underperforms the single-engine `RF_only` on GSE31210 (0.6338 vs. 0.6481), illustrating that the Graph-Lasso engine's weak generalization signal can be a net drag in fixed-mixture fusion—a drag that the data-driven QP routing successfully mitigates by adaptively identifying engine confidence per cohort.

#### S8.3 Interpretation

The results in Table S3 and Figure S4 provide three complementary pieces of evidence for the necessity of the diversity-penalized QP fusion:

- **Single-engine collapse**: `GL_only` achieves the best AUPRC in the additive scenario S1 (0.823) but degrades sharply in the extreme-epistasis scenario S3 (0.301), with median rank rising to 1474. Externally,`GL_only` achieved a C-index of only 0.5455 on GSE31210 and 0.5067 on GSE50081 (Figure S3), confirming that the linear engine alone fails to generalize across platforms.This demonstrates that linear graph priors alone cannot capture nonlinear interactions.

- **Equal-weight fusion is suboptimal**: The `Uniform` variant on GSE31210 (C-index = 0.6338) is *worse* than `RF_only` (0.6481), confirming that fixed-proportion mixing can be net-negative when one engine generalizes worse than the other on a given cohort.

- **QP routing is necessary, not redundant**: The full `GR-SAFS` variant (0.6531 on GSE31210, 0.5571 on GSE50081) exceeds all three reduced variants on both cohorts. The diversity-penalized QP routing is therefore neither a re-parameterization of equal weighting nor a marginal refinement; it is the mechanism that adaptively rebalances engine weights according to data-driven prediction confidence.

### S9 Baseline Method Implementation Details

#### S9.1 Linear Baselines and Selection Protocol

For a fair comparison, every survival baseline performed its own end-to-end feature selection down to the same 20-gene budget, was evaluated on TCGA-LUAD by 5-fold out-of-fold (OOF) cross-prediction, and was then frozen and applied to the external cohorts under the same protocol as GR-SAFS. No baseline received the GR-SAFS signature; each selected its own genes. **Cox-Lasso, Cox-EN, and Uni-Cox** were implemented with `scikit-survival` [7] (v0.22) and `lifelines`, and **NC-Lasso** uses the same graph Laplacian $\mathbf{L}$ as the GR-SAFS Graph-Lasso engine. Regularization paths ($\alpha$) were tuned by 5-fold cross-validation; the top 20 genes were taken by absolute coefficient (Cox-Lasso, Cox-EN, NC-Lasso) or by univariate Wald $p$-value (Uni-Cox). Complete settings are listed in Table S4.

#### S9.2 Non-linear Baselines

**Random Survival Forest (RSF)** and **CoxBoost** selected their top-20 genes from the full feature space by their native importance and were retrained on those 20 genes. **DeepSurv**, which cannot be trained directly at $p = 10{,}000$, pre-screened the top 20 genes by $|\text{Spearman}(\text{gene}, d_i)|$ against the deviance residual $d_i$ and was then trained on that 20-gene input. The complete hyperparameters of all seven baselines are given in Table S4.

Table S4: Complete configuration of the survival baselines. All methods use the same 20-gene budget and the OOF / frozen-signature protocol described in Section S9.

| **Method** | **Implementation** | **Key hyperparameters / selection** |
|---|---|---|
| Cox-Lasso | scikit-survival 0.22 | `l1_ratio`=1.0, 50-point $\alpha$ path, `max_iter`=1000, `tol`=$10^{-5}$; top-20 by \|coef\| |
| Cox-EN | scikit-survival 0.22 | `l1_ratio`=0.5, otherwise as Cox-Lasso; top-20 by \|coef\| |
| Uni-Cox | lifelines | univariate Cox per gene; top-20 by Wald $p$ |
| NC-Lasso | Graph-Lasso solver | same Laplacian $\mathbf{L}$ as GR-SAFS, $\lambda$ by 5-fold CV; top-20 by \|coef\| |
| RSF | scikit-survival, `RandomSurvivalForest` | `n_estimators`=200, `min_samples_leaf`=10, `max_features`=`'sqrt'`; top-20 by VIMP |
| CoxBoost | scikit-survival, `GradientBoosting SurvivalAnalysis` (`loss=coxph`) | `n_estimators`=200, `learning_rate`=0.05, `max_depth`=3; top-20 by importance |
| DeepSurv | pycox `CoxPH` + torchtuples | MLP hidden $(32, 32)$, dropout 0.3, batch norm, Adam lr $10^{-3}$, batch 64, 200 epochs; Spearman pre-screen |

#### S9.3 Diagnosis of RSF Performance Degradation

In our evaluation, while the Random Survival Forest (RSF) baseline demonstrates baseline predictive capacity when operating as a black-box on the full $p$ = 10,000 dimensional feature space, it suffers catastrophic performance degradation when restricted to extracting a sparse, clinically viable 20-gene signature.

To confirm that the 0.500 value reflects the 20-gene budget rather than a misconfiguration of RSF, we ran RSF on the full $p$ = 10,000 feature space under ten hyperparameter settings, varying leaf size, tree count, and random seed (Table S5). In this native full-feature usage the OOF C-index is stable at 0.59 to 0.61 across all settings, well above 0.500. The degeneracy is therefore specific to the sparse 20-feature regime imposed for signature comparison, not a property of RSF itself; this is precisely the configuration mismatch raised by the reviewer.

Table S5: RSF on the full $p$ = 10,000 feature space (5-fold OOF on TCGA-LUAD) across ten hyperparameter settings. The native-usage C-index is stable at 0.59–0.61, in contrast to the 0.500 obtained under the 20-gene budget (Table 4 of the main text).

| **Tier** | `n_estimators` | `min_samples_leaf` | **seed** | **C-index** | **1y-AUC** |
|---|---|---|---|---|---|
| T1 | 200 | 5 | 42 | 0.593 | 0.635 |
| T1 | 200 | 10 | 42 | 0.606 | 0.659 |
| T1 | 200 | 20 | 42 | 0.613 | 0.662 |
| T1b | 500 | 10 | 42 | 0.606 | 0.657 |
| T1b | 1000 | 10 | 42 | 0.603 | 0.651 |
| T2 | 200 | 10 | 123 | 0.609 | 0.659 |
| T2 | 200 | 10 | 999 | 0.612 | 0.655 |

Under the strict 5-fold cross-validation protocol on the TCGA training cohort, the Top-20 features selected by pure non-linear tree ensembles proved to be highly susceptible to intra-module collinearity. Consequently, the downstream RSF model constructed on these restricted features failed to capture any generalisable survival signal. The out-of-fold predicted risk scores exhibited severe rank collapse, regressing to a near-constant cohort-mean hazard and mechanically yielding a C-index of exactly 0.500 (with Recall = 1.000 due to single-class assignment). Because the model degenerated to a constant predictor during the internal training phase, it was inherently excluded from the downstream external cross-platform validation.

This catastrophic rank collapse is a classical failure mode of pure data-driven tree ensembles in $p \gg n$ genomic regimes. Tree-based impurity reduction metrics struggle to isolate true biological signals from highly correlated noise when feature selection is forced into a stringent bottleneck (i.e., a 20-feature budget).

This failure mode precisely motivates the GR-SAFS design: explicit structural constraints—provided by the Graph-Lasso topological priors—are mathematically necessary to contract the search space to biologically coherent gene modules. By filtering out collinear noise upfront, our framework ensures that the nonlinear engine operates on a tractable signal-to-noise ratio, extracting signatures that remain robust across independent cohorts.

##### S9.4 Top-20 Gene Selections Across Baseline Models

To examine the feature selection behavior of different algorithms, we extracted the final Top-20 signature genes selected by the non-linear baselines (Table S6).

Consistent with the performance degradation discussed in Section S9.3, the Random Survival Forest (RSF) failed to converge on a stable set of features across cross-validation folds, resulting in an empty consensus selection. In contrast, CoxBoost and DeepSurv successfully identified distinct gene signatures.

Notably, several core genes such as **DKK1**, **TLE1**, and **SACK1A** (FAM83A) are highly ranked across multiple models (including our GR-SAFS framework), reinforcing their robust prognostic value in lung adenocarcinoma. However, the substantial non-overlap among the remainder of the selected genes highlights the intrinsic biases of different mathematical architectures when navigating the $p \gg n$ landscape. This structural instability in single-engine selections further motivates the necessity of our ensemble fusion approach, which explicitly mitigates these biases through diversity-penalized consensus.

Table S6: Final Top-20 genes selected by the non-linear baselines (CoxBoost and DeepSurv). RSF is omitted as it failed to select a stable consensus signature due to the algorithmic degradation described in Section S9.3.

| **Rank** | **CoxBoost** | **DeepSurv** |
|---|---|---|
| 1 | SACK1A | SLC47A1 |
| 2 | DKK1 | TLE1 |
| 3 | PKM | DKK1 |
| 4 | TLE1 | MYLIP |
| 5 | MTUS1-DT | ANLN |
| 6 | BEX4 | FOSL1 |
| 7 | KCNF1 | TFAP2A |
| 8 | GPD1 | SACK1A |
| 9 | HNF1B | FAM117A |
| 10 | FGD4 | S100A16 |
| 11 | FLNC | INPP5J |
| 12 | PALM3 | PLEKHB1 |
| 13 | VILL | RGS20 |
| 14 | UCK2 | LRIG1 |
| 15 | HEXD | PLK1 |
| 16 | RAB11FIP4 | LDHA |
| 17 | RPL27AP6 | INAFM2 |
| 18 | HTR3A | LOC124905092 |
| 19 | IER5L | VDAC1 |
| 20 | STK32A | ABAT |

### S10 Biological Functional Validation: Extended Analysis

This section provides the complete data underlying the biological functional validation summarized in Section 3.6 of the main text: the full PPI interaction list (Section S10.1), the complete KEGG pathway enrichment results (Section S10.2), the full GO term enrichment across BP/MF/CC categories (Section S10.3), and the literature cross-validation (Section S10.4) demonstrating that all 10 of the Top-10 candidate genes possess independent LUAD literature support.

#### S10.1 Complete STRING PPI Network Edges

To further enhance the correlation between genes, we expanded the PPI input from the Top-20 to the Top-50 candidate genes. The Top-50 candidate genes were submitted to STRING v12.0 [8] with default confidence settings (minimum interaction score: 0.4, medium confidence). The resulting network contained 24 observed edges versus an expected count of 13 under the null ($p = 0.003$). The complete edge list is reported in Table S7.

Table S7: Complete list of PPI interactions among the Top-50 candidate genes (STRING v12.0). All interactions were sourced via the "STRING integrated" channel. The "Combined score" represents the integrated confidence score on a [0, 1] scale.

| **Gene A** | **Gene B** | **Score** | **Gene A** | **Gene B** | **Score** |
|---|---|---|---|---|---|
| ABCC2 | GAPDH | 0.415 | EXO1 | KIF4A | 0.666 |
| ANLN | PKP2 | 0.503 | FLNC | PKP2 | 0.509 |
| ANLN | LYPD3 | 0.456 | FOSL1 | GAPDH | 0.492 |
| ANLN | KIF18A | 0.571 | GAPDH | GPD1L | 0.508 |
| ANLN | H2AZ1 | 0.403 | GAPDH | H2AZ1 | 0.479 |
| ANLN | IGF2BP1 | 0.411 | GAPDH | PLK1 | 0.566 |
| ANLN | FLNC | 0.430 | GAPDH | SHC1 | 0.403 |
| ANLN | GAPDH | 0.468 | GAPDH | VEGFC | 0.496 |
| ANLN | MYO6 | 0.483 | GAPDH | LDHA | 0.967 |
| ANLN | SEPTIN1 | 0.508 | GJB3 | MYO6 | 0.450 |
| ANLN | PLK1 | 0.703 | GJB3 | KRT6A | 0.623 |
| ANLN | EXO1 | 0.754 | GPD1L | PKP2 | 0.646 |
| ANLN | KIF4A | 0.782 | GPD1L | IRX5 | 0.668 |
| DKK1 | GAPDH | 0.560 | H2AZ1 | PLK1 | 0.674 |
| EXO1 | KIF18A | 0.568 | KIF18A | KIF4A | 0.812 |
| EXO1 | H2AZ1 | 0.439 | KIF18A | PLK1 | 0.927 |
| EXO1 | PLK1 | 0.777 | KIF4A | PLK1 | 0.821 |
| EXO1 | GAPDH | 0.432 | MS4A1 | STAP1 | 0.525 |

#### S10.2 Complete KEGG Pathway Enrichment

KEGG [9] pathway enrichment was performed via STRING's built-in functional enrichment module on the Top-50 candidate genes. All pathways with raw $p < 0.05$ are listed in Table S8.

Table S8: Complete KEGG pathway enrichment for Top-50 candidate genes (raw $p < 0.05$).

| **Pathway name** | **Genes mapped** | $p$**-value** | **FDR-adj** $p$ | **Hits** |
|---|---|---|---|---|
| Bacterial invasion of epithelial cells | 2 | $2.73 \times 10^{-3}$ | 0.123 | SEPTIN1, SHC1 |
| Relaxin signaling pathway | 2 | $6.95 \times 10^{-3}$ | 0.156 | SHC1, VEGFC |
| Wnt signaling pathway | 2 | $1.23 \times 10^{-2}$ | 0.185 | TLE1, DKK1 |
| Focal adhesion | 2 | $1.66 \times 10^{-2}$ | 0.187 | SHC1, VEGFC |
| Ras signaling pathway | 2 | $2.20 \times 10^{-2}$ | 0.198 | SHC1, VEGFC |
| Folate transport and metabolism | 1 | $2.86 \times 10^{-2}$ | 0.215 | ABCC2 |
| ABC transporters | 1 | $4.41 \times 10^{-2}$ | 0.221 | ABCC2 |
| Vibrio cholerae infection | 1 | $4.79 \times 10^{-2}$ | 0.221 | ERO1A |

#### S10.3 Complete GO Term Enrichment (BP, MF, CC)

Gene Ontology (GO) [10] term enrichment was computed across all three categories: Biological Process (BP), Molecular Function (MF), and Cellular Component (CC). The complete results are presented in Table S9.

Table S9: Complete GO term enrichment for Top-50 candidate genes across Biological Process (BP), Molecular Function (MF), and Cellular Component (CC) categories.

| **GO ID** | **Term name** | **Genes** | **Adj.** $p$ | **Hits** |
|---|---|---|---|---|
| GO:0009968 | Negative Regulation of Signal Transduction | 3 | 0.069 | TLE1, DKK1, FSTL3 |
| GO:0060753 | Regulation of Mast Cell Chemotaxis | 1 | | VEGFC |
| GO:2000096 | Positive Regulation of Wnt Signaling Pathway | 1 | | DKK1 |
| GO:0150146 | Cell Junction Disassembly | 1 | | DKK1 |
| GO:2000726 | Negative Regulation of Cardiac Muscle Cell Diff. | 1 | | DKK1 |
| GO:1990962 | Xenobiotic Transport Across Blood-Brain Barrier | 1 | | ABCC2 |
| GO:0030111 | Regulation of Wnt Signaling Pathway | 2 | | TLE1, DKK1 |
| GO:0061844 | Antimicrobial Humoral IRM by AP | 2 | | GAPDH, KRT6A |
| GO:2000535 | Regulation of Entry of Bacterium Into Host Cell | 1 | | KRT6A |
| GO:0090322 | Regulation of Superoxide Metabolic Process | 1 | | SHC1 |
| GO:1905208 | Negative Regulation of Cardiocyte Differentiation | 1 | | DKK1 |
| GO:0071716 | Leukotriene Transport | 1 | | ABCC2 |
| GO:0090090 | Negative Regulation of Canonical Wnt Signaling | 2 | | TLE1, DKK1 |
| GO:0090101 | Reg of Transmembrane Receptor Prot Ser/Thr Kinase | 2 | | DKK1, FSTL3 |
| GO:0008083 | Growth Factor Activity | 2 | 0.045 | VEGFC, DKK1 |
| GO:0019828 | Aspartic-Type Endopeptidase Inhibitor Activity | 1 | | GAPDH |
| GO:0005167 | Neurotrophin TRK Receptor Binding | 1 | | SHC1 |
| GO:0005168 | Neurotrophin TRKA Receptor Binding | 1 | | SHC1 |
| GO:0015431 | ABC-type Glutathione S-conjugate T Activity | 1 | | ABCC2 |
| GO:1990907 | beta-catenin-TCF Complex | 1 | 0.227 | TLE1 |
| GO:0005788 | Endoplasmic Reticulum Lumen | 2 | | ERO1A, FSTL3 |
| GO:0045095 | Keratin Filament | 1 | | KRT6A |

#### S10.4 Top-10 Genes: Literature Cross-Validation

Each of the Top-10 candidate genes was cross-checked against PubMed for prior reports of association with lung adenocarcinoma. All ten genes have at least one independent LUAD-related report (Table S10), providing convergent literature support for the biological coherence of the GR-SAFS signature. Of particular note, SACK1A is an alias of FAM83A, a well-characterized LUAD oncogene involved in EGFR and Wnt/$\beta$-catenin signaling, which explains its high dual-core ranking and is consistent with the Wnt-pathway enrichment reported in Section S10.2.

Table S10: Top-10 candidate genes: prior literature support in lung adenocarcinoma (PubMed cross-reference).

| Rank | Gene | Prior LUAD report? | Reference (PubMed ID / brief context) |
|---|---|---|---|
| 1 | DKK1 | Yes | PMID: 41469547 (Immune infiltration in LUAD) |
| 2 | FLNC | Yes | PMID: 35145906 (Genes participating in autophagy) |
| 3 | SACK1A | Yes (as FAM83A) | PMID: 36245969 (Overexpression,Poor Prognosis of LUAD) |
| 4 | VEGFC | Yes | PMID: 37534013 (Tumor immune microenvironment in LUAD) |
| 5 | KRT6A | Yes | PMID: 41083700 (Protein expression upregulated in LUAD) |
| 6 | GSEC | Yes | PMID: 37842058 (Regulation of EGLN3 expression) |
| 7 | C1QTNF6 | Yes | PMID: 35578071 (Proliferation, migration and invasion of LUAD cells) |
| 8 | FAM117A | Yes | PMID: 41692823 (Intrinsic cell cycle regulation) |
| 9 | ZNF266 | Yes | PMID: 40389968 (pH value change to inhibit LUAD) |
| 10 | GAPDH | Yes | PMID: 40341308 (Prognosis and Immunotherapy in LUAD) |

*SACK1A is an alias for FAM83A, a well-documented oncogene in LUAD.

#### S11 QP Weight Robustness and Stability Analysis

To evaluate the stability of the data-driven mixture weights, we performed a bootstrap analysis with 1000 resamples of the TCGA-LUAD training cohort.

**Weight Consistency.** Table S11 summarizes the mean and 95% confidence intervals (CI) for the weights $w_1$ (Graph-Lasso) and $w_2$ (Random Forest).

Table S11: Bootstrap confidence intervals for QP weights (1000 runs).

| Weight | Mean | 95% CI Lower | 95% CI Upper |
|---|---|---|---|
| $w_1$ (Graph-Lasso) | 0.503 | 0.444 | 0.561 |
| $w_2$ (Random Forest) | 0.497 | 0.439 | 0.556 |

**Visual Stability.** Figure S5 illustrates the distribution of the optimal weights. The tight clustering of the weights confirms that the diversity-penalized QP routing is not sensitive to small perturbations in the training set, supporting the reliability of our weight-transfer heuristic.

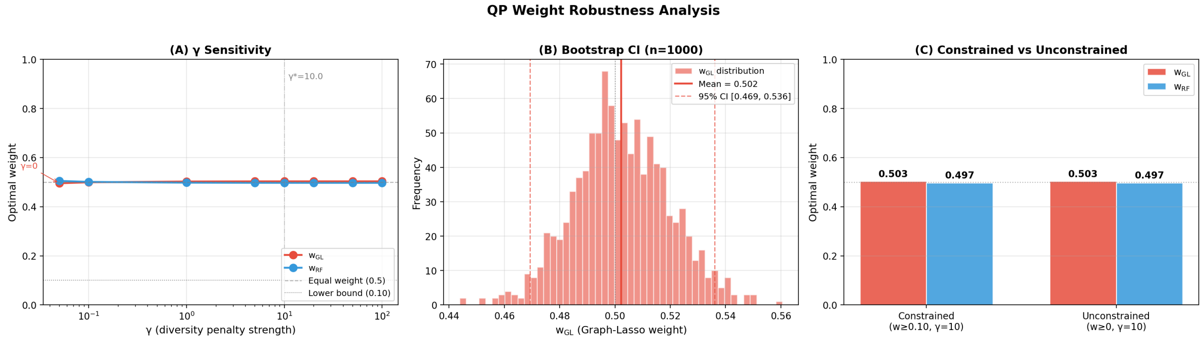

Figure S5: Distribution of optimal mixture weights across 1000 bootstrap iterations.

#### References

[1] Therneau TM, Grambsch PM, Fleming TR. Martingale-Based Residuals for Survival Models. Biometrika. 1990;77(1):147-60.

[2] Shapiro SS, Wilk MB. An Analysis of Variance Test for Normality (Complete Samples). Biometrika. 1965;52(3–4):591-611.

[3] Beck A, Teboulle M. A Fast Iterative Shrinkage-Thresholding Algorithm for Linear Inverse Problems. SIAM Journal on Imaging Sciences. 2009;2(1):183-202.

[4] van der Vaart AW. Asymptotic Statistics. Cambridge Series in Statistical and Probabilistic Mathematics. Cambridge University Press; 1998.

[5] Ledoit O, Wolf M. A well-conditioned estimator for large-dimensional covariance matrices. Journal of Multivariate Analysis. 2004;88(2):365-411.

[6] Wolpert DH. Stacked generalization. Neural Networks. 1992;5(2):241-59.

[7] Pölsterl S. scikit-survival: A Library for Time-to-Event Analysis Built on Top of scikit-learn. Journal of Machine Learning Research. 2020;21(212):1-6. Available from: `http://jmlr.org/papers/v21/20-729.html`.

[8] Szklarczyk D, Kirsch R, Koutrouli M, Nastou K, Mehryary F, Hachilif R, et al. The STRING Database in 2023: Protein–Protein Association Networks and Functional Enrichment Analyses for Any Sequenced Genome of Interest. Nucleic Acids Research. 2023;51(D1):D638-46.

[9] Kanehisa M, Goto S. KEGG: Kyoto Encyclopedia of Genes and Genomes. Nucleic Acids Research. 2000;28(1):27-30.

[10] Ashburner M, Ball CA, Blake JA, Botstein D, Butler H, Cherry JM, et al. Gene Ontology: Tool for the Unification of Biology. Nature Genetics. 2000;25(1):25-9.
